# Towards the implementation and interpretation of masked ICA for identifying signatures of autonomic activation in the brainstem with resting-state BOLD fMRI

**DOI:** 10.1101/2024.12.20.628885

**Authors:** Mary Miedema, Kyle T.S. Pattinson, Georgios D. Mitsis

## Abstract

The brainstem is the site of key exchanges between the autonomic and central nervous systems but has historically presented a challenging target for study with BOLD fMRI. A potentially powerful although under-characterized approach to identifying nucleic activation within the brainstem is masked independent component analysis (mICA), which restricts signal decomposition to the brainstem itself, thus aiming to reduce the strong effect of physiological noise in nearby regions such as ventricles and large arteries. In this study, we systematically investigate the use of mICA to uncover signatures of autonomic activation in the brainstem at rest. We apply mICA on 40 subjects in a high-resolution resting state 7T dataset following different strategies for dimensionality selection, denoising, and component classification. We show that among the noise mitigation techniques investigated, cerebrospinal fluid denoising makes the largest impact in terms of mICA outcomes. We further demonstrate that across preprocessing pipelines and previously reported results the majority of components are spatially reproducible, but temporal outcomes differ widely depending on denoising strategy. Evaluating both hand-labelling and whole-brain specificity criteria, we develop an intuitive framework for mICA classifications. Finally, we make a comparison between mICA and atlas-based segmentations of brainstem nuclei, finding little consistency between these two approaches. Based on our evaluation of the effects of methodology on mICA and its relationship to other signals of interest in the brainstem, we provide recommendations for future uses of mICA to identify autonomically-relevant BOLD fluctuations in subcortical structures.

## Introduction

The autonomic nervous system (ANS), which includes the parasympathetic and sympathetic branches, is well-known to modulate systemic physiological activity such as cardiac and respiratory regulation^1^. Task-based functional imaging studies have shown that a group of regions in the brain including the brainstem, thalamus, hypothalamus, amygdala, insula, cingulate cortex as well as the prefrontal cortex - collectively termed the central autonomic network (CAN) - are associated with the processing of autonomic function^2^. Moreover, several recent studies^3–7^ have used resting-state fMRI data to identify regions in the brain associated with but not limited to the CAN which are strongly coupled with autonomic control of the cardiovascular system. At the mechanistic core of such investigations lies the brainstem, linking the cortical systems of the central nervous system through complex interchanges to regulatory branches of the ANS.

Despite a growing interest in understanding the neural correlates of brain-body interactions, a major difficulty in interpreting brainstem data remains that the blood-oxygen-level-dependent (BOLD) contrast measured with fMRI is an indirect measure of neuronal activity, where associated changes in BOLD are generated by an underlying hemodynamic response. BOLD measurements are confounded by physiological noise such as non-neuronal fluctuations in signal driven by changes in respiratory and cardiac activity as well as blood gas levels, which often cause synchronous signal modulations that can strongly influence functional connectivity analysis^8–13^. In an autonomic context, changes in physiology drive both neural and mechanical processes detectable with BOLD, which can be considered both signal and noise, respectively (**Figure 1**).

**Figure 1:**
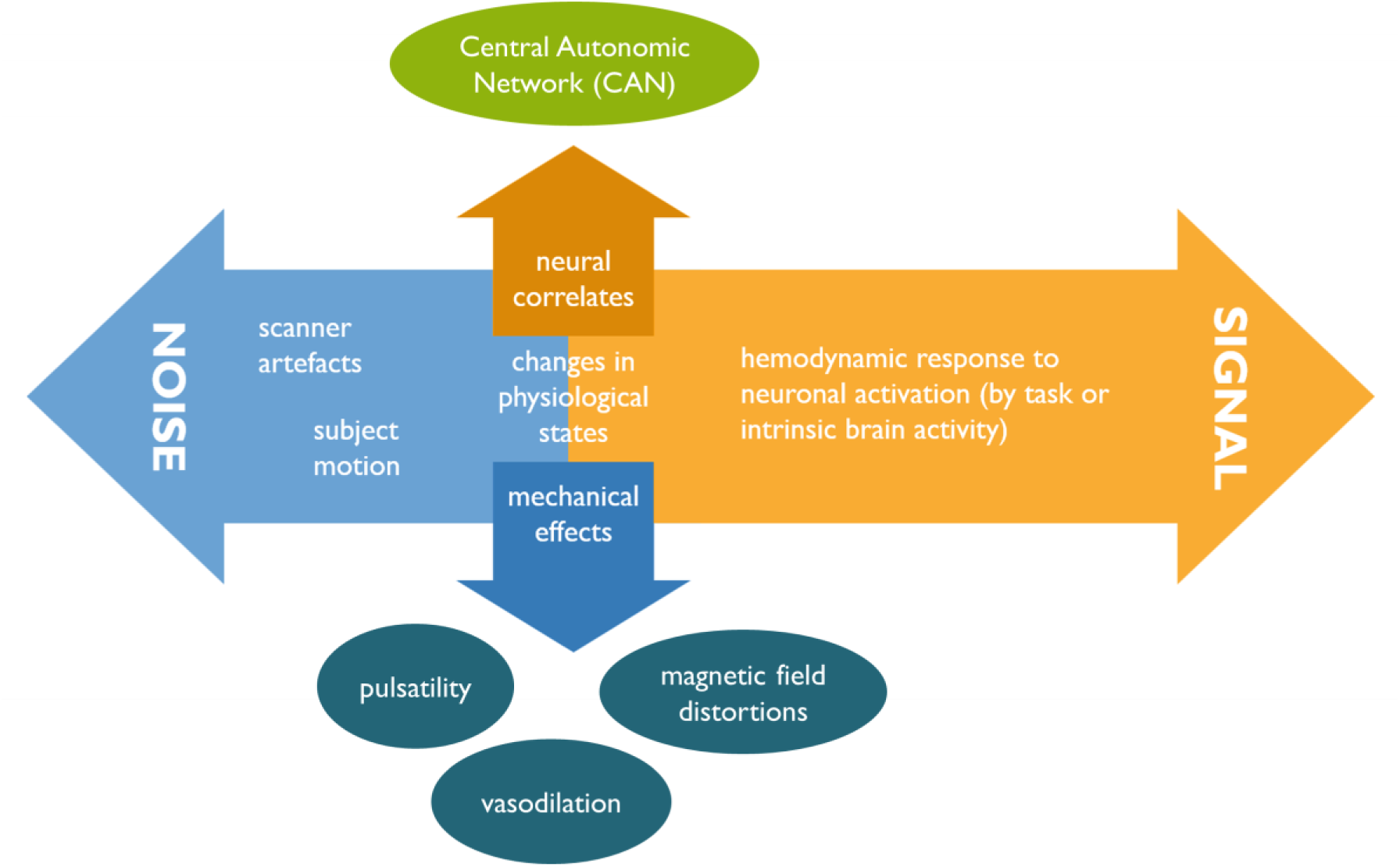
Autonomic activity can affect changes in BOLD measurements through both neuronal and non-neuronal pathways. For fMRI investigations concerned with the involvement of autonomic activity in cognitive processes, changes in physiological state during a scan can be associated with both signal and noise. While the central autonomic network (CAN) is implicated most directly in the neural integration of autonomic activity, other networks may likewise exhibit changes in the activation or strength of neural pathways corresponding to changes in physiological state^16^. However, changes in physiological state will necessarily reflect the contribution of non-neuronal i.e. mechanically-driven processes to the BOLD signal. These include but are not limited to the pulsatile motion of fluid driven by cardiac activity, shifts in bulk susceptibility causing distortions of the magnetic field during changes in breathing, or vasodilatory responses to concentrations of carbon dioxide in the bloodstream,^14^ which may all be treated as artefactual confounds to any subsequent neuronal interpretation of the fMRI data.

The problem of identifying such autonomically-driven signal is particularly compounded in the brainstem, where proximity to strong sources of noise -- such as nearby cerebrospinal fluid and large blood vessels as well as the movement of the chest -- significantly increases the proportion of non-neuronal confounds contaminating BOLD fluctuations^14^. Nonetheless, accurately separating neuronally-relevant signal fluctuations in the brainstem from noise remains crucial in interpreting fMRI data with respect to brain-body interactions. Regions of the brainstem associated with autonomic control may interact with established cortical networks^15^, resulting in a neuronal modulation of the BOLD signal entangled with undesirable artefacts caused by changes in systemic physiology.

In recent years, numerous studies addressing physiological noise in BOLD fMRI have reported functional connections between the brainstem and regions of the brain associated with autonomic control under resting-state conditions^17^. Such investigations often rely on anatomical atlases for delineating seeds to obtain functional connectivity patterns^18–20^, which are subject to pitfalls in the robust identification of the small centres of activation inherent to brainstem nuclei combined with the challenge of accurate group-level registration in subcortical regions^21^. Moreover, since the brainstem is highly susceptible to physiological noise contamination, the appropriate noise correction methods prior to signal extraction remain an issue of some contention^22^. In contrast to the application of *a priori* brainstem atlases or model-based de-noising techniques, independent component analysis (ICA) has been proposed as a more sensitive method of spatially differentiating noise from signal.^23^ Specifically, masked independent component analysis (mICA), first proposed and demonstrated in the brainstem by Beissner et al.^24^, has been suggested^25–27^ to provide the best solution for a data-driven decomposition of meaningful signals within subcortical structures without contamination from nearby noise sources such as the cerebrospinal fluid (CSF).

Despite its singularity as the only brainstem-specific method for data-driven de-noising in recent fMRI literature, implementation of mICA in the brainstem remains somewhat rare, although several studies have recently demonstrated promising mICA decompositions in brainstem regions during scan conditions involving explicit autonomic modulation^28–30^ or have otherwise attempted mICA decompositions in the resting-state^31,32^. A brief overview of these fMRI studies which have used mICA to isolate neuronally-relevant signal in the brainstem is presented in Table 4 and discussed in detail in the following sections. Critically interpreting the application and consistency of mICA across these studies is challenging: wide differences exist between the preprocessing strategies employed prior to mICA application, while the spatial and temporal characteristics of the mICA decomposition are infrequently reported amongst study results. The implementation of mICA requires many study-specific choices such as those illustrated in Figure 2, which may be difficult to navigate and have significant effects on study outcomes. While in their original paper^24^ Beissner et al. found that applying physiological noise modelling^33^ prior to mICA did not significantly change their results, the effects of alternative noise correction strategies have not been likewise evaluated. Moreover, although mICA was originally proposed and validated on resting-state fMRI data^24^, subsequent focus on task-based autonomic activation has obscured how and if masked ICA can be used to isolate meaningful functional activity in the brainstem during resting-state conditions. To-date, reliable functional data-driven segmentation of resting-state brainstem BOLD signals corresponding to autonomic regulation has not been achieved using mICA.^34^

**Figure 2:**
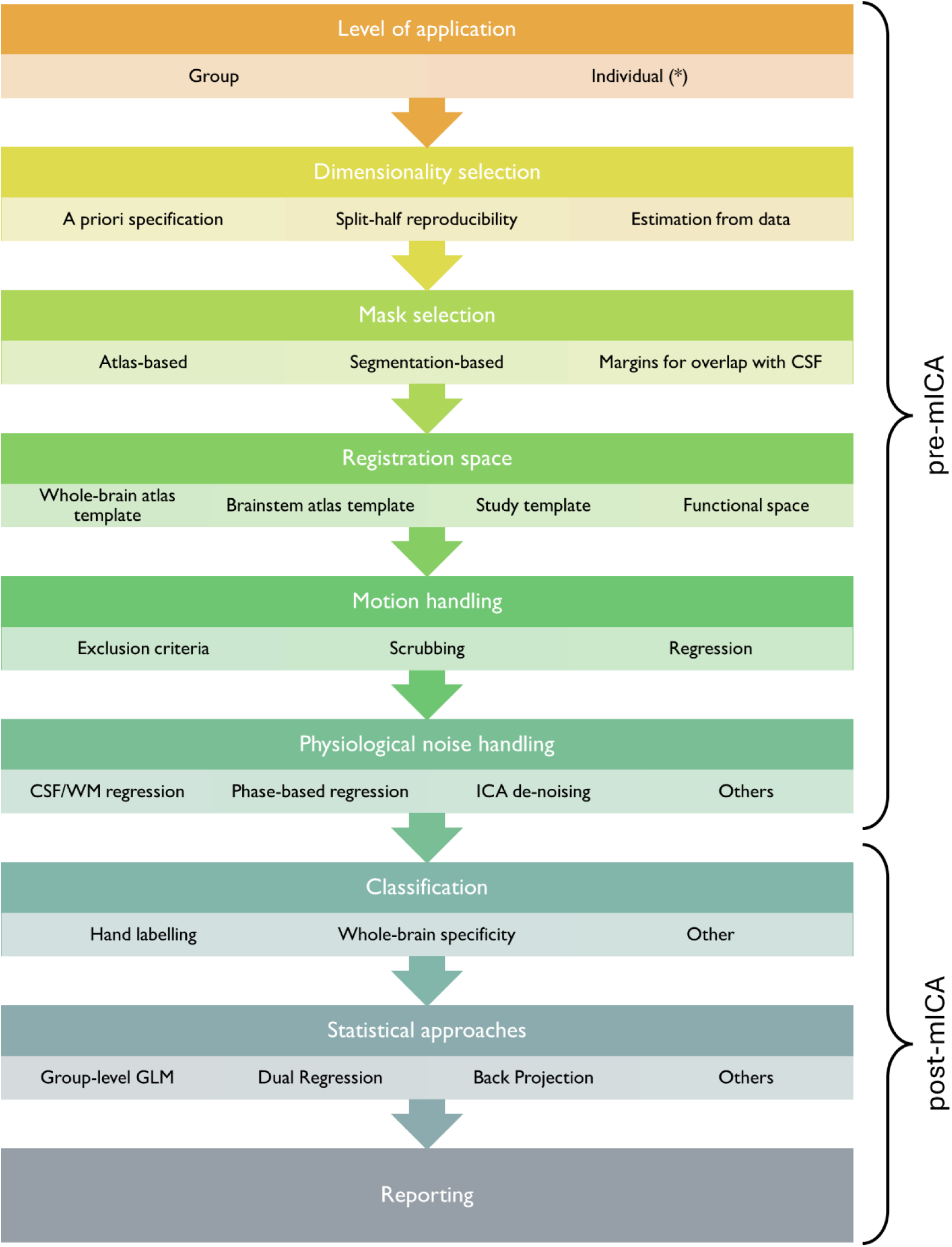
The application of masked independent component analysis (mICA) necessitates numerous decisions regarding the preprocessing applied to functional data prior to mICA, the selection of parameters such as dimensionality and mask for mICA itself, and the usage and interpretation of mICA results. Here some of the different possibilities for each stage of analysis are shown, as extrapolated from current fMRI studies using mICA (see Table 4). *: the application of mICA on the single-subject level is uncommon and presents its own particular challenges; see the corresponding Discussion section.

Overall, the particular challenges posed by physiology and anatomy in the brainstem position flexible data-driven approaches as a promising solution for identifying signals relevant to brain-body interactions in this crucial structure. However, without clear guidelines for their implementation and interpretation, tools such as mICA may remain only tangential to the course of ongoing brainstem investigations. The present work therefore sets out to characterize the effect of different pre-processing strategies in the context of detecting autonomic-related activity resting-state brainstem BOLD fMRI data using mICA. We demonstrate that the application of mICA is highly influenced by the approach to dimensionality estimation of the underlying data but is robust to several different combinations of d physiological noise correction. Highlighting the classification of components as autonomically relevant signal or noise, we propose several strategies for evaluating and reporting the results. Crucially, our work addresses previously unanswered questions in the practical usage of mICA for the brainstem in resting-state datasets . To this end, we employ a high-resolution (7T) dataset and we perform a comparison with atlas-based segmentations of brainstem nuclei, as well as investigate the relation between mICA outputs (components) to systemic low-frequency oscillations linked to autonomic regulation.

## Materials and methods

### Data acquisition

We used high-resolution resting-state data from 40 healthy subjects previously acquired in a study on cortical connectivity associated with respiratory control using a 7 T Siemens Magnetom scanner^35^. For this study, ethics approval was given by the Oxfordshire Clinical Research Ethics and subjects gave written, informed consent prior to participation. For each participant, we used 9.5 minutes of eyes-open resting-state data with a T2*-weighted, gradient echo planar image (EPI) sequence (TE = 24 ms; TR = 3 s (190 volumes); flip angle, 90 °; FOV, 220 mm; 2 mm isotropic resolution) along with T1-weighted MPRAGE structural scans (TE = 2.96 ms; TR = 2200 ms; flip angle, 7 °; FOV, 224 mm; 0.7 mm isotropic resolution), field maps, and cardiac and respiratory physiological signals (500 Hz).

### Physiological data processing

Cardiac and respiratory signals were processed using Matlab (v.R2023a) with a combination of custom scripts and the PhysIO toolbox^36^. Following bandpass filtering and peak detection, variance between interbeat intervals were used to calculate heart-rate variability (HRV) using a sliding-window approach. Likewise, extracted respiratory peaks and amplitudes were used to calculate changes in respiration volume per time (RVT). The calculated HRV and RVT were then convolved with their respective canonical physiological response functions^37,38^. Cardiac and respiratory phase time series for each subject were also extracted for use as described in the subsequent de-noising section.

### Pre-processing of functional and structural data

#### Initial pre-processing

Rs-fMRI data pre-processing was carried out using FEAT (FMRI Expert Analysis Tool) Version 6.04, which is part of FSL^39^ (FMRIB’s Software Library, www.fmrib.ox.ac.uk/fsl). Initial pre-processing steps included deleting the first 3 volumes of each fMRI series to allow the magnetic field to reach a steady state, followed by motion correction, distortion correction^40^ using the fieldmap images through FUGUE, and highpass temporal filtering with a cutoff frequency of 0.01 Hz. The functional images were co-registered to the brain-extracted high-resolution T1-weighted images using boundary-based registration^41^. At this stage, no smoothing was applied to the functional data. The functional data were then registered to structural images and the MNI152 (Montreal Neurological Institute) atlas using FNIRT (nonlinear registration with FMRIB’s Nonlinear Image Registration Tool) and resampled to 1 mm isotropic resolution. Finally, the structural data were segmented into CSF, grey matter (GM) and white matter (WM) compartments using FEAT (FMRIB’s Automated Segmentation Tool).^42^

#### Overview of de-noising pipelines

The minimally pre-processed data were then fit to a general linear model (GLM) with a design matrix corresponding to nuisance regressors to account for signal drift, subject motion, and physiological sources of noise according to one of eight pre-processing pipelines. These pipelines are described in detail in Table 2. Briefly, pipelines were designed to encompass a range of pre-processing strategies, from minimally to very aggressive in terms of removing variance associated with noise. Two different strategies were employed to account for physiological noise by including RETROICOR^43^ and CSF-extracted regressors^44^. In the latter case, principal component analysis (PCA) was performed on CSF voxels lying within a sphere 80 mm in diameter centred on the brainstem and components explaining 50% of the variance in this region were included as regressors in the noise model^45^.

**Table 1:**
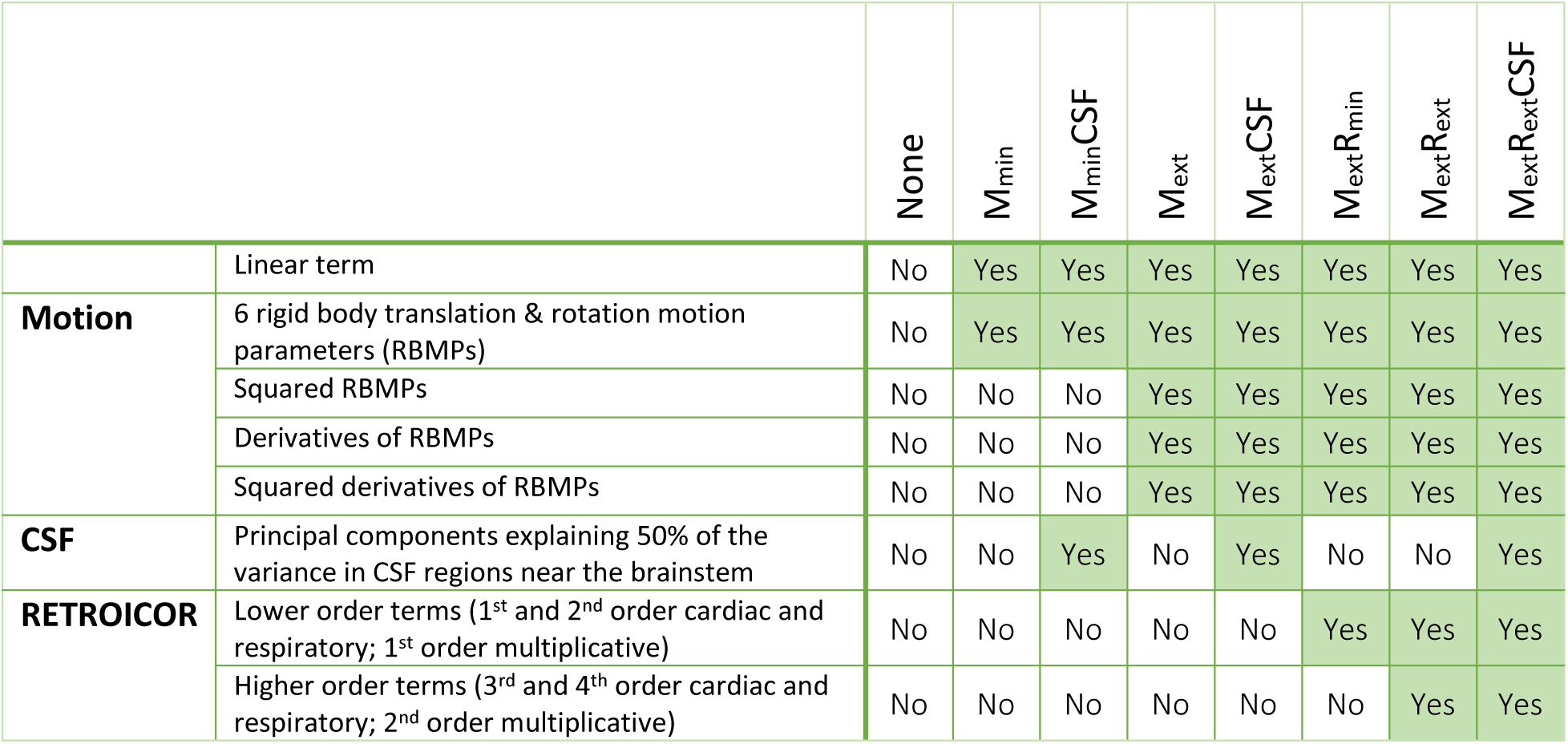
Parameters included in a general linear model (GLM) to de-noise functional data within the brainstem for each of the 8 examined pipelines. All terms were mean-subtracted and normalized before inclusion in the GLM.

**Table 2:** Median dimensionalities estimated on single-subject data for each pipeline, used in subsequent group mICA.

| Pipeline | NoCI | M<sub>min</sub> | M<sub>min</sub>CSF | M<sub>ext</sub> | M<sub>ext</sub>CSF | M<sub>ext</sub>R<sub>min</sub> | M<sub>ext</sub>R<sub>ext</sub> | M<sub>ext</sub>R<sub>ext</sub>CSF |
| --- | --- | --- | --- | --- | --- | --- | --- | --- |
| **Median dimensionality** | 32 | 30 | 30 | 30 | 29 | 29 | 30 | 27 |

### Application of masked independent component analysis

#### Mask delineation

All masked analyses used the anatomical brainstem mask provided in the supplementary materials of

Beissner et al.’s foundational paper on mICA^24^ in the resting-state for more useful comparisons between the former and the present study. This mask was defined based on the ICBM152 template and thresholded conservatively with manual removal of non-brainstem regions to minimize noise introduced into the masked volume by voxels corresponding to CSF.

#### Dimensionality selection

In its initial implementation, decomposition dimensionality for group mICA was estimated by first performing mICA at the single-subject level with automatic dimensionality estimation, and subsequently using the median dimensionality value to decompose the group data.^24^ Thus, to facilitate direct comparisons, single-subject pre-processed data in MNI152 space were analyzed separately for each pipeline using probabilistic ICA^46^ with automatic dimensionality estimation implemented in MELODIC. Values ranging from 27-32 estimated by MELODIC were then used in subsequent group mICA as the so-termed median dimensionality for each pipeline (see Results; Table 2).

In more recent implementations of mICA, notably the mICA toolbox^25^, split-half resampling of the data is recommended to assess spatial reproducibility of mICs across a pre-specified range of potential dimensionalities. However, this technique has largely been used for task datasets, typically resulting in reproducible task-originating spatial activation patterns, and thus requires further validation in resting-state data. Using the mICA toolbox (v.1.18), the 40 subjects were sorted into 50 randomly-selected split-half subgroups, over which mICA was applied for dimensionality values ranging from 5-40. For each set of subgroups and values, the components were sorted to optimize spatial similarity using the toolbox’s implementation of the Hungarian sorting algorithm^47^ for maximizing the average spatial correlation between components across the set. From this analysis (see Results; Figure 3), a value of 8 was chosen as the so-termed reproducible dimensionality for all pipelines.

**Figure 3:**
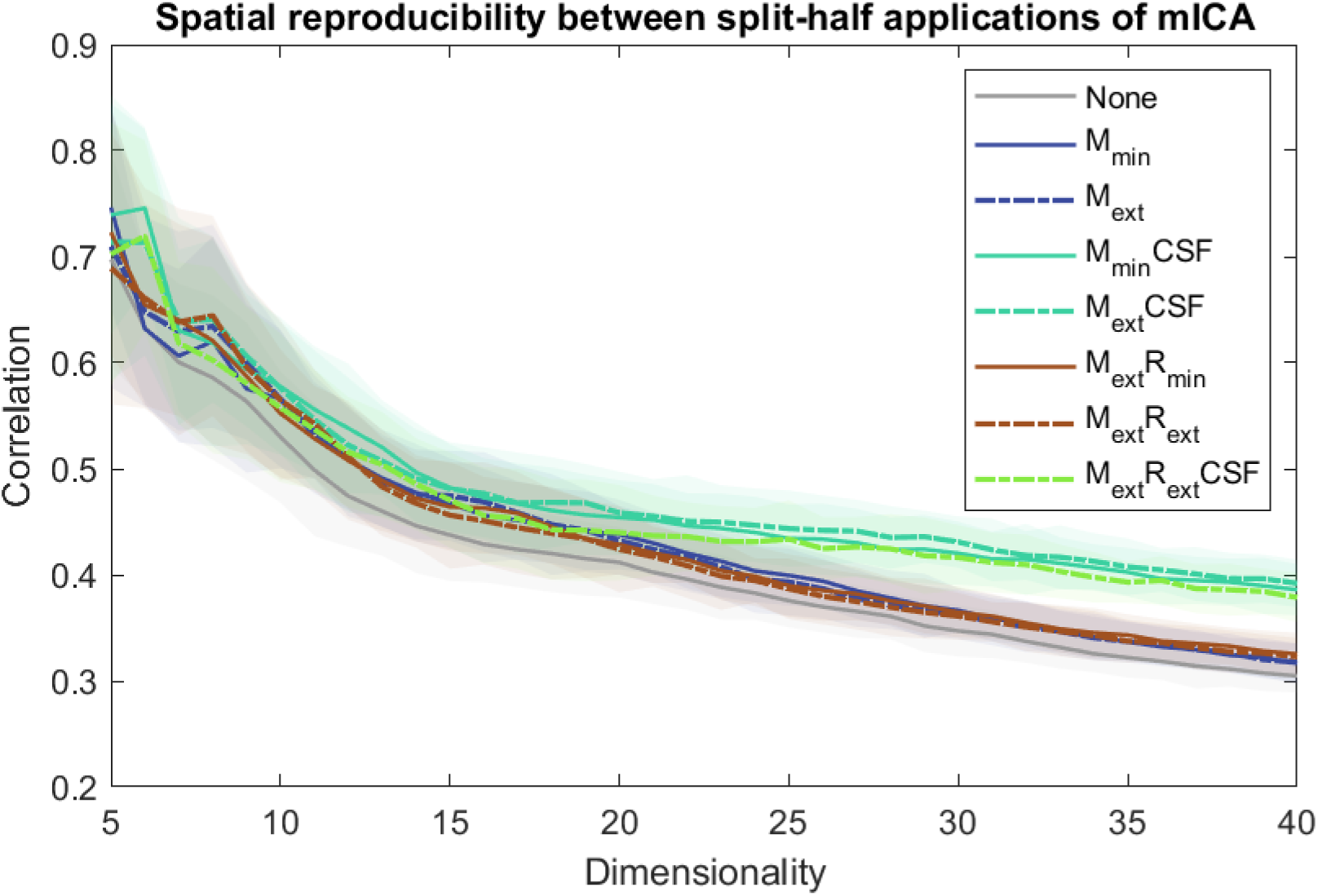
Spatial reproducibility across split-half subgroups declines with higher dimensionality values chosen for mICA decomposition; for higher dimensionalities CSF de-noising increases reproducibility. Spatial similarity here is calculated as the average spatial correlation between matched component pairs across split-half datasets (50 repetitions) to which mICA was applied for a range of pre-specified dimensionalities for each de-noising pipeline. Shaded regions indicate 95% confidence intervals.

#### Decomposition with mICA

After de-noising with each pipeline, the pre-processed data were transformed into MNI152 space and underwent voxelwise normalization. Data across subjects were then temporally concatenated and analyzed using a group ICA^46^ to obtain mICA independent components (mICs) for both median and reproducible dimensionalities within the masked brainstem region. A Gaussian mixture-modelling technique was used to threshold the estimated component maps.

#### Hand classification of mICA components

Each mIC was labelled as either ‘signal’, ‘noise’, or ‘ambiguous’ based on its spatial and temporal characteristics in accordance with established guidelines for whole-brain ICA^48^. Briefly, a focal spatial distribution was considered characteristic of signal, while spatial distributions near or following the brainstem’s edges (i.e. overlapping with CSF regions) were considered characteristic of noise. Time courses for a given mIC were obtained for each subject using dual regression^49^ and plotted for each subject, along the average power spectra across subjects and the power spectrum of the mIC amplitude from the group decomposition. These temporal characteristics were used to further identify characteristics of noise such as sudden jumps in time courses and frequency peaks below 0.01 Hz. Due to the relatively long TR of the fMRI dataset, in which many higher frequency signatures of noise may be aliased into lower frequencies, spatial characteristics were considered more reliable than temporal characteristics for assessing mIC quality. A label of ‘signal’, ‘noise’, or ‘ambiguous’ was therefore assigned first based on spatial characteristics. Temporal characteristics were then used to further label spatially ‘ambiguous’ mICs. Whole-brain connectivity maps (as described in the following section) were not included for consideration during labelling.

### Spatial replication analysis

The reproducibility of the mICs estimated for each dimensionality and pipeline with the 37 original mICs previously reported in 7T resting-state data by Beissner et al.^24^ was assessed in an analysis analogous to that performed for the reproducible dimensionality estimation described previously. It should however be noted that, while reproducibility has been typically reported as the spatial correlation between unthresholded mICs, here, correlations were estimated between the openly available thresholded original mICs and this study’s unthresholded mICs.

### Component-based connectivity analysis

#### Creation of whole-brain connectivity maps

After group mICA was performed on data for each pipeline and both dimensionalities, the time courses estimated for each mIC were extracted and de-meaned for each subject. A brain-wide PCA was conducted to obtain a set of brain-wide suitable regressors and whole-brain functional data was cleaned using the regressors corresponding to each pipeline (Table 1). Finally, the thus-denoised whole-brain data was smoothed with a 5 mm kernel. Connectivity to the smoothed whole brain data for each pipeline was evaluated using a general linear model across subjects. Parameter estimates for each pipeline, dimensionality, and mIC were evaluated for statistical significance using a nonparametric permutation test implemented in Randomise^50^ with 5000 permutations and family-wise error correction to *p < 0.01* to obtain maps of whole-brain connectivity at the group level.

#### Specificity determination

Previously, assigning either signal or noise labels to individual mICs has been performed by calculating a specificity measure from each mIC’s whole-brain connectivity profile. Originally, this measure was defined as one minus the ratio of activation within the CSF compartment divided by activations within the white and gray matter^24^. In one particular implementation, another version of this measure was generated within only a restricted subcortical volume instead of the whole brain.^29^ Here, we propose a modified definition of specificity defined as the product of thresholded whole-brain connectivity maps (*A*) and thresholded compartmental probability maps (*M*), normalized by an additional term reflecting activation in the CSF compartment:

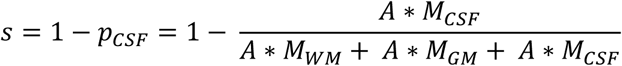

This modified equation includes normalization to improve interpretability by avoiding the calculation of negative specificity values in instances of high connectivity to CSF compartments. We also propose an additional measure of specificity to assess the presence of co-fluctuations within the central autonomic network (CAN) using a mask of regions associated with autonomic control:

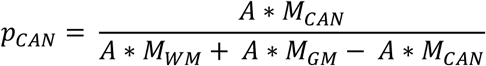

Here, the mask associated with activation of the CAN (*M*_*CAN*_) was generated from a compilation of spheres 6 mm in diameter centered at the MNI coordinates reported for nineteen clusters associated with sympathetic and parasympathetic regulation in a meta-analysis of autonomic function^2^.

### Atlas-based connectivity analysis

Structural regions of interest from the Harvard Ascending Arousal Network Atlas^51^ corresponding to two nuclei strongly linked to autonomic arousal -- the locus coeruleus (LC) and periaqueductal gray (PAG) -- were transformed into functional space. For each subject, the mean signal in each region was extracted, temporally de-meaned, and fit to whole-brain de-noised data corresponding to each pipeline with a general linear model. Similar to mIC connectivity, parameter estimates for each pipeline and nucleus were evaluated for statistical significance using a nonparametric permutation test implemented in Randomise^50^ with 5000 permutations and family-wise error correction to *p < 0.01* to obtain maps of whole-brain connectivity across subjects.

## Results

### Dimensionality and spatial reproducibility of mICA within pipelines

Overall, dimensionality estimation and reproducibility scores were remarkably consistent across pipelines. The median dimensionality estimated by MELODIC on single-subject data ranged from 27-32, with lower dimensionalities tending to correspond to less de-noised data (Table 2), suggesting that the removal of nuisance regressors gradually reduced spatial variability in the brainstem data.

However, the split-half reproducibility investigation demonstrated that higher dimensionalities reduced the spatial similarity of components identified across subgroups of the dataset (Figure 3). In general, spatial similarity scores were fairly low (<0.8), although scores classified as reproducible by Beissner et al.^24^ (>0.5) were achieved for dimensionalities ranging from 5-12 for most pipelines. However, at very low dimensionalities (5, 6, and 7 for some pipelines), the MELODIC algorithm did not converge to a solution for several split-half subgroups. Thus, a dimensionality of 8 was chosen to represent the most reproducible application of group mICA across pipelines.

It should be noted that for higher dimensionalities, such as the median values selected for group mICA by estimation of single-subject data, Figure 3 shows that pipelines incorporating CSF denoising exhibited higher reproducibility scores than the other pipelines. This could suggest greater inter-subject variability in the spatial distribution of pulsatile noise components, perhaps directly linked to anatomical variability.

### Classification of mICA components

Figure 4 shows specificity curves across all pipelines for both dimensionality implementations and their connectivity profiles calculated using either smoothed or unsmoothed whole-brain data. Notably, unlike the sole previous specificity curve reported for masked ICA in the brainstem^24^, no clear plateau - below which non-specific components may be rigorously classified - exists. As well, we obtained notably lower specificity values than those previously reported for both smoothed and unsmoothed data, even using the modified norm (Eq. 1) which would be expected to inflate the observed specificity values. Using smoothed whole-brain data tended to increase specificity in less-specific components, presumably due to the combined effect of reducing inter-subject variability and blurring connectivity across compartments. In contrast, the lower specificity values obtained without smoothing exhibited a less linear curve, which could be used to threshold noise components in a more straightforward manner.

**Figure 4:**
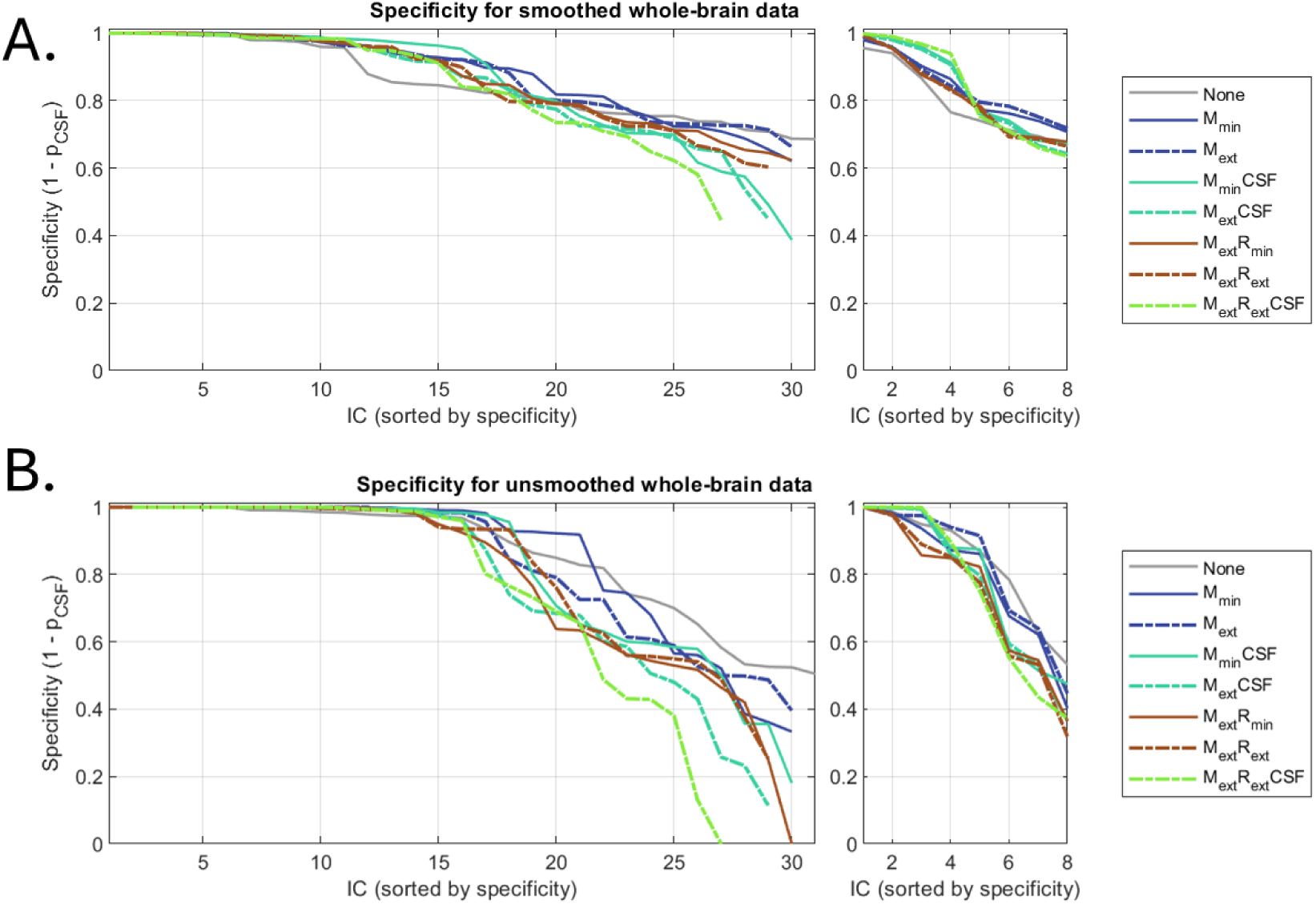
The specificity of whole-brain connectivity exhibits a more dynamic range across components when calculated over unsmoothed data. Specificities across pipelines with median dimensionality were calculated separately using smoothed whole-brain data (A) and unsmoothed whole-brain data (B). Left panels show high (median) dimensionality applications of mICA, while right panels show low (reproducible) dimensionality mICA. A lower specificity indicates a whole-brain connectivity profile predominating in the CSF compartment, which suggests that it should be classified as noise.

Without consideration of its calculated specificity value, each mIC was individually classified according to its spatial and temporal characteristics within the brainstem as either signal, noise, or ambiguous. Analogously to whole-brain ICA, spatial characteristics such as focality or proximity to the edges of the brainstem mask (where motion of the ventricles introduces significant pulsatility artefacts) can inform manual classification of components.^48^ Figure 5 gives an example of some representative components and their classifications for components derived from the un-denoised (None) pipeline.

**Figure 5:**
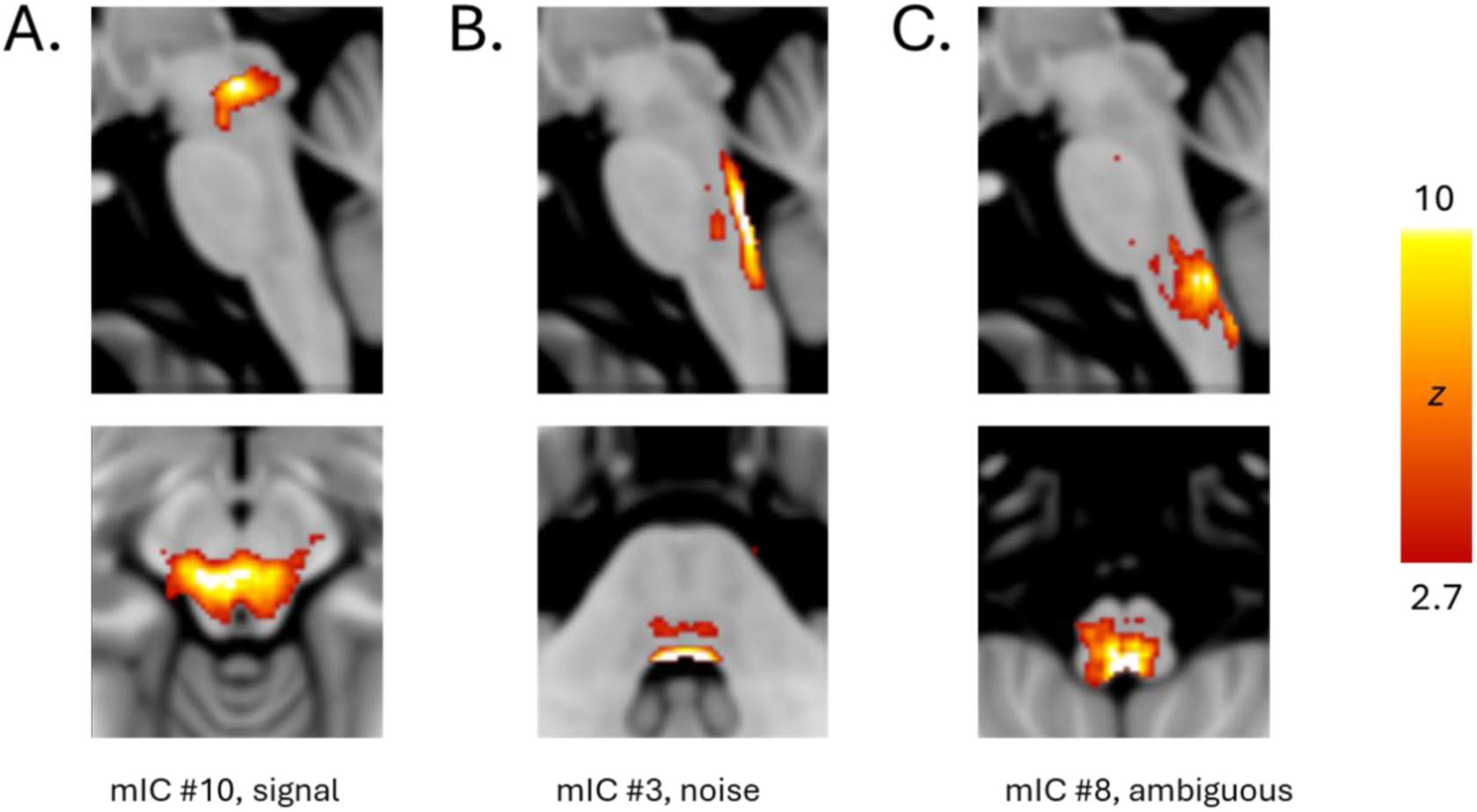
Some exemplar classifications (taken from the None pipeline) showing the thresholded spatial maps of mICs manually labelled as signal (A), noise (B), or too ambiguous to be reliably assigned (C). Here, the single focal peak of component 10 located centrally in the midbrain is suggestive of signal, while the dispersed peak along the fourth ventricle of component 3 is suggestive of noise. Of particular note is the doubled spatial distribution of component 3, where a secondary activation was often found anterior to the primary boundary activation of similar components across pipelines. In contrast, component 8 has a more dispersed peak near the boundary of the medulla, a region in which physiological noise effects are quite strong. The interpretation of this component is ambiguous. In general, hand-labelling of mICS relied heavily on their spatial characteristics, since the TR (3 s) of the data and the group-level application of mICA made temporal interpretations challenging, although frequency spectra and dual regression time courses for each mIC were used to supplement classification.

After classification, the specificity scores for each category of hand-labelled component were assessed for both unsmoothed (Figure 6) and smoothed (Supplementary Figure 1) whole-brain data. This analysis was conducted only for the median dimensionality instance for each pipeline, since the granularity of the higher dimensional mICA provided a larger number of noise and signal components for comparison and generally better separated noise from signal. Broadly, good agreement between whole-brain specificity and hand-labelling of components as signal or noise validates the former metric for classification. Moreover, Figure 6 demonstrates that increased denoising improves the separation of noise and signal specificity scores, pointing towards the possibility of a specificity threshold which could be selected for automated classification. For example, setting a specificity threshold of 0.9 would give a true positive rate of 0.981 and false positive rate of 0.075 across all pipelines (excluding ambiguous components). The range of specificity scores within the ambiguous category of components also highlights that this more objective criterion could be useful where hand-labelling falls short.

**Figure 6:**
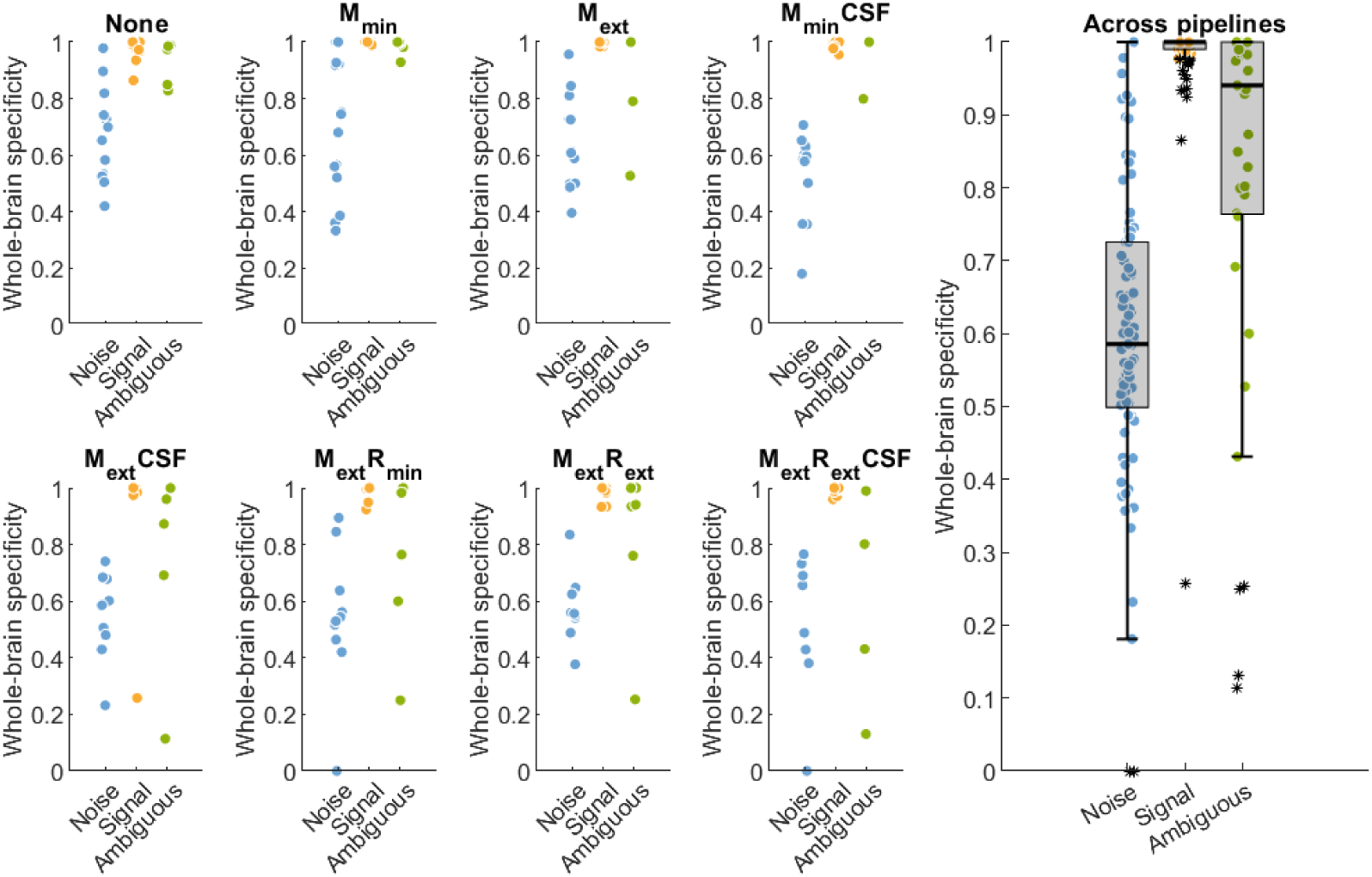
Hand-labelling of individual components shows moderate agreement with corresponding specificity values (calculated using unsmoothed whole-brain data). Results for each individual pipeline are shown on the left; their aggregate is on the right. The inclusion of motion correction parameters in the noise regression model seems to improve the specificity scores of signal components, while the implementation of physiological noise correction (whether RETROICOR or CSF-derived) reduces the specificity scores of noise components.

### Spatial reproducibility of mICA with previously reported findings

To assess the reproducibility of components between this study and those resulting from Beissner et al.’s original implementation of mICA^24^, we followed an analogous procedure to the split-half reproducibility comparison. However, we chose a minimum threshold of 0.4 for spatial correlation to reflect reproducibility (vs. 0.5 in Beissner et al.’s original analysis), since their components provided were thresholded (reducing the correlations). The two studies under comparison had highly similar data (7T resting-state acquired with a TR of 2.52 and 3s, isotropic resolution of 2.5 and 2 mm, and 50 and 40 subjects for the original study^24^ and the present one, respectively). However, where the original study performed only phase-based physiological noise modelling and reported insignificant differences in a comparative implementation without physiological noise correction, we tested a wide range of denoising combinations, as well as the effect of the reproducible dimensionality selection recommended in the subsequent benchmarking paper for mICA^25^. Figure 7 demonstrates that the lower reproducible dimensionality implementation in general poorly replicates the 37 original components, as expected due to the low specificity of these components. In contrast, the higher median dimensionality implementations matched reproducibly with a subset of 20 of the original components across all pipelines. However, different denoising combinations affected which components were reproducibly found across pipelines: only 8 original components were reproducibly matched by a majority of pipelines (Figure 7, lower left). Moreover, it should be noted that while 32 out of the 37 total components were classified as specific (i.e. signal) in the original study, the specificity scores and hand-labelling of components in the present study suggest a much more dominant role was played by noise in our data and resulting mICs, even for more aggressive denoising approaches (Figure 6).

**Figure 7:**
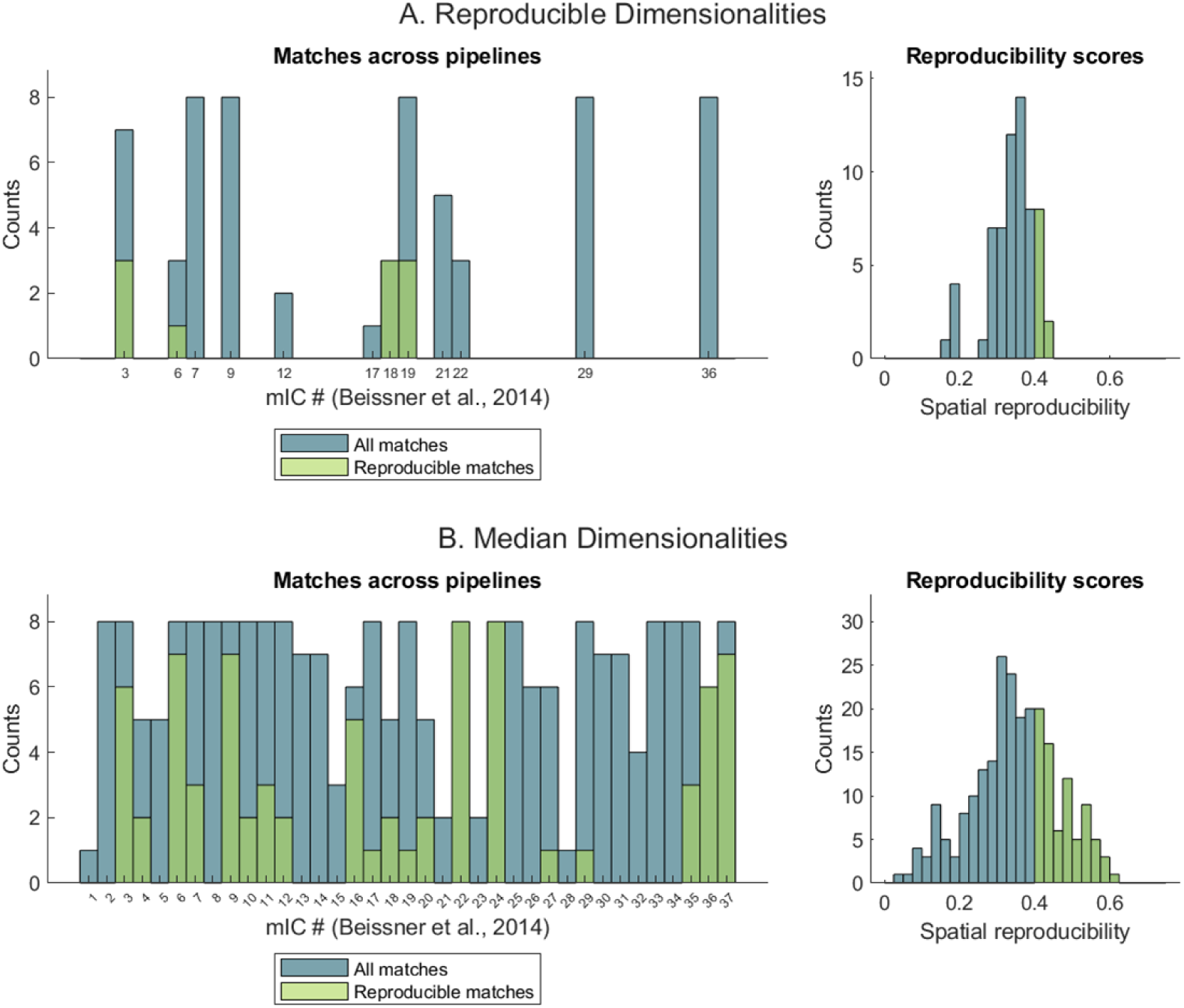
Spatial reproducibility scores between mICS identified by this study and Beissner et al.’s original resting-state mICA results (2014), demonstrating the effect of dimensionality selection and pipeline on replicability. Eleven components (out of 37 possible) were selected as matches for the eight mICs in the reproducible dimensionality case (A), while all 37 components were selected at least once across all the possible pipelines in the median dimensionality mICA application (B). Of these matched components, few achieved a reproducible (>0.4) score (right), although matches that achieved reproducibility tended to be reproducible across more than one denoising pipeline (left).

The distribution of reproducibility scores for matched components did not vary significantly between pipelines (Supplementary Figure 2). A full list and representation of all reproducibly-matched components can be found in Supplementary Figure 3.

### Effects of dimensionality selection on mICA outcomes

Since the identification of mIC sources, whether noise or signal, is generally purposed for use in a general linear model or other time-based analysis to assess connectivity with broader regions of the brain, we wished to assess the temporal similarity of mICs within pipelines based on the dimensionality selected for decomposition. Figure 8 demonstrates the correlation between time-series of mICs within pipelines and across both the median and reproducible dimensionalities, along with physiologically-relevant and atlas-extracted signals for further comparison (see later discussion). The higher dimensionality of the median approach led to splitting of some of the mICs identified using the lower, reproducible dimensionality across all pipelines, as is particularly clear for mICs explaining larger proportions of variance in the brainstem. For instance, the first ‘reproducible’ mIC is in general strongly correlated to the first one to three ‘median’ mICs within each pipeline. Figure 8 also suggests that increased denoising tends to reduce the temporal anti-correlation between mICs, both intra- and inter-dimensionally. This temporal independence within mICs is advantageous for later statistical validity in constructing a design matrix of higher rank or indeed for efficacy of identifying separate sources of BOLD signal fluctuations.

**Figure 8:**
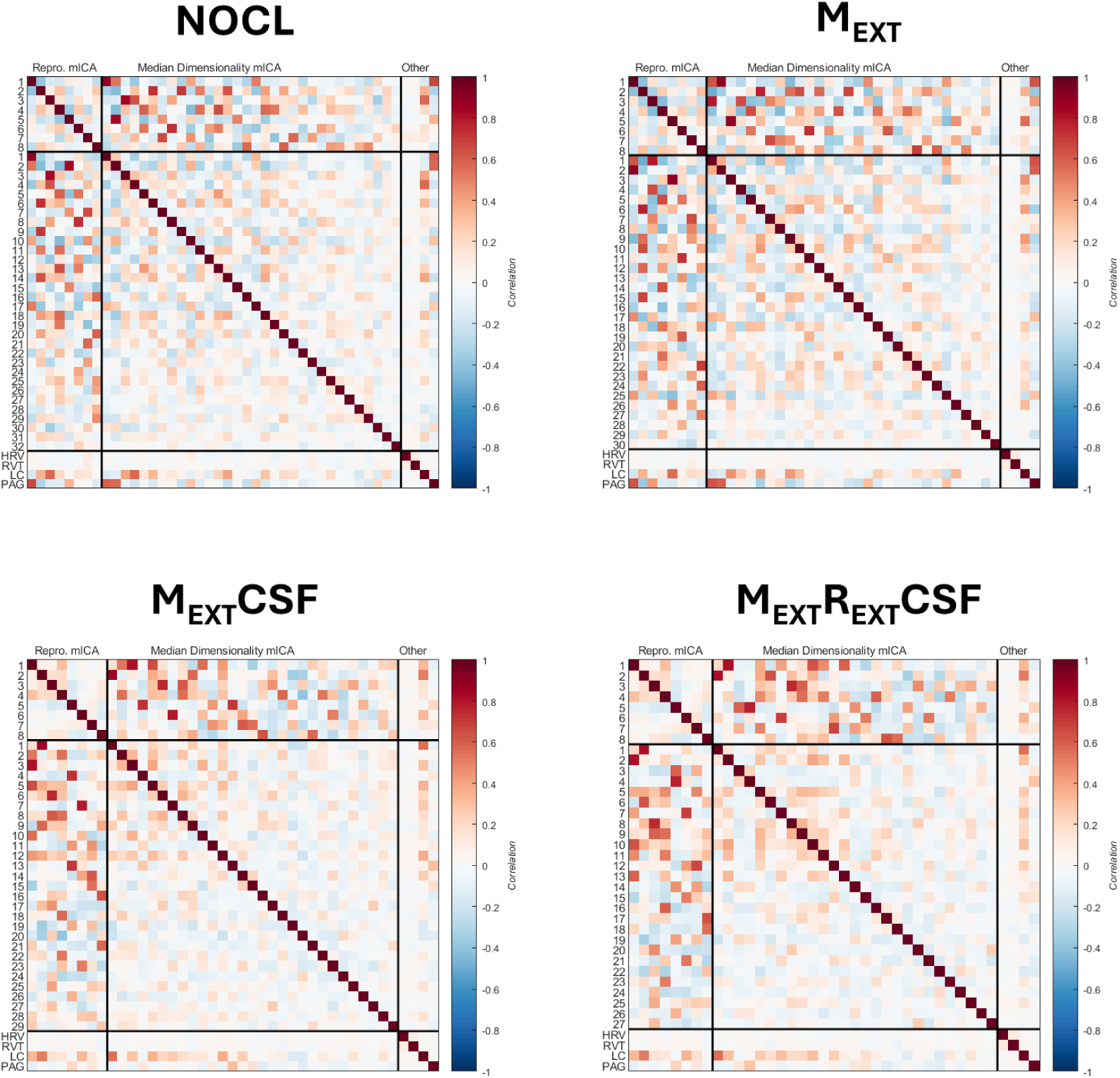
Higher dimensionality mIC implementation splits temporal variance across several components. Results are presented across several representative pipelines. The temporal correlation between mICs within pipelines is calculated between the reproducible and median dimensionalities specified for decomposition and averaged across subjects. Correlation with physiological signals of interest (HRV, RVT) and ROI-extracted signals (LC, PAG) are provided for comparison (last four rows/columns).

### Effects of denoising on mICA outcomes

We further wished to interrogate whether mICs may be used to isolate physiological noise similarly to the other denoising techniques we applied. Thus, the temporal correlation between regressors generated for our de-noising pipelines and the mICs obtained for the un-denoised data (‘None’ pipeline) was calculated (Figure 9). Weak correlations between mICs and regressors such as motion in the x-direction suggest that some mICs are more linked than others to conventional denoising techniques. However, overall, little temporal similarity between regressors and mICs was observed. The mICs explaining higher amounts of variance in the un-denoised data yielded higher correlation values with the PCA-extracted signals from CSF near the brainstem.

**Figure 9:**
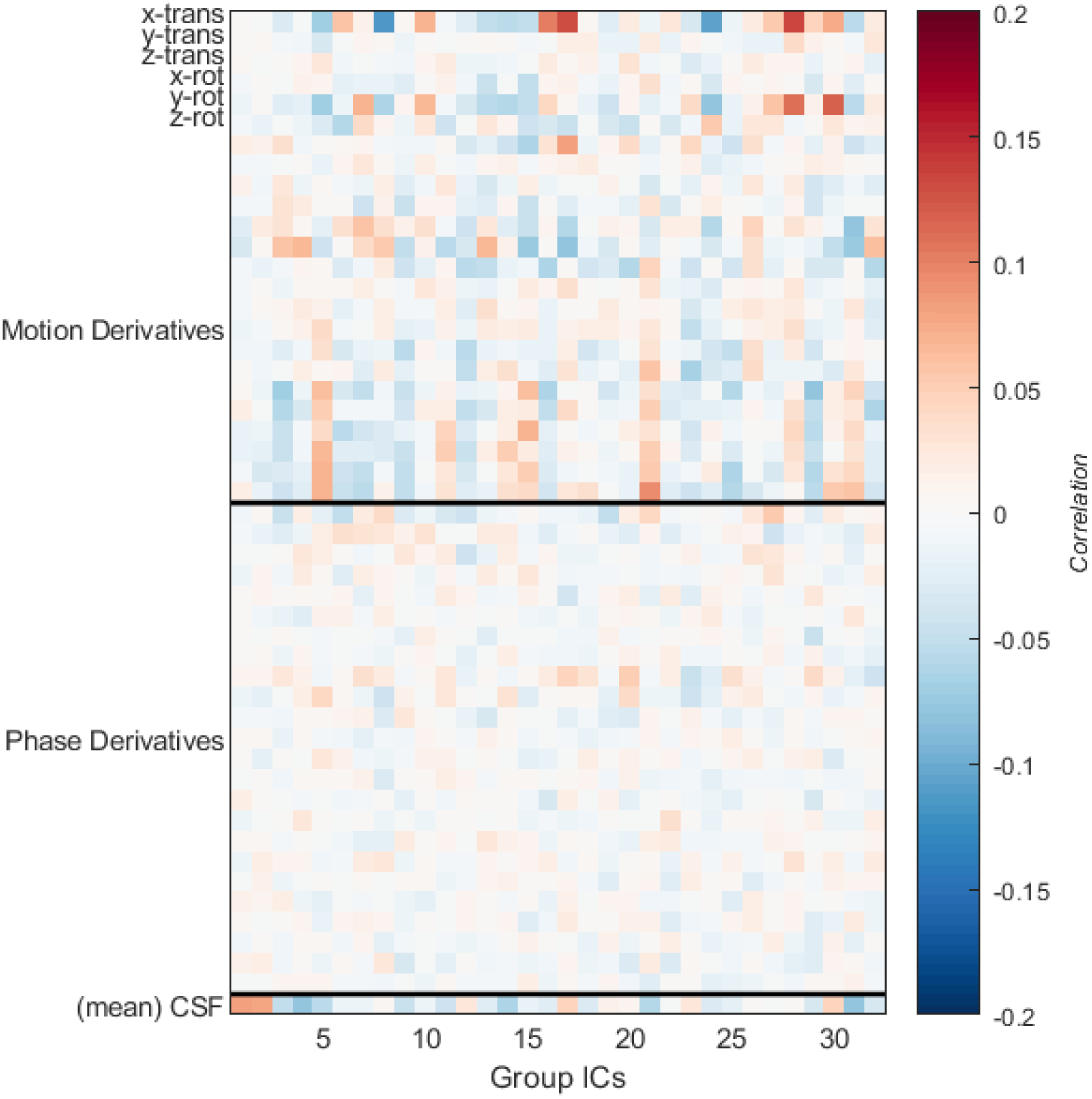
Regardless of classification as noise or signal, mICs obtained without prior denoising isolate different temporal dynamics than physiological noise models. Here the temporal correlation between regressors used in the employed denoising pipelines (see Table 2) and the mIC time-courses calculated for the ‘None’ pipeline and median dimensionality are averaged across all subjects. The mean of the correlation value over all CSF regressors is plotted because the number of principal components extracted from the CSF compartment varied for each subject.

Exploring the distribution of these characteristics across subjects, Figure 10 suggests that all mICs were more strongly correlated with CSF regressors than with motion or physiological phase-derived regressors. This is particularly notable, as previous applications of mICA did not typically include CSF de-noising techniques (Table 1), and the benchmark introduction of mICA in the brainstem examined only the effect of phase-based physiological denoising on mICA application.^24^ In this study, Beissner et al. suggested that the effect of physiological denoising was minimal on mICA outcomes, reporting similar spatial decompositions and reproducibility scores of mIC components derived with and without phase-based cardiac and respiratory denoising. However, our results suggest that temporal outcomes should also be considered, and that different approaches to de-noising, particularly CSF-based, should be considered. Indeed, while the primary motivation for applying a mask in this analysis is to avoid introducing proximal confounding CSF fluctuations to BOLD measurements originating within the brainstem, our results suggest that the mICA approach exhibits only limited success in this regard. To highlight the difference in possible outcomes, Figure 11 shows that for the least and most aggressively denoised data (the ‘None’ and ‘M_ext_R_ext_CSF’ pipelines, respectively) to which mICA was applied, the temporal correlation between mICs was typically low and – where existent – was split across several mICs within the two different pipelines. A full exploration of temporal similarity of mICA outcomes between pipelines is provided in Supplementary Figure 4.

**Figure 10:**
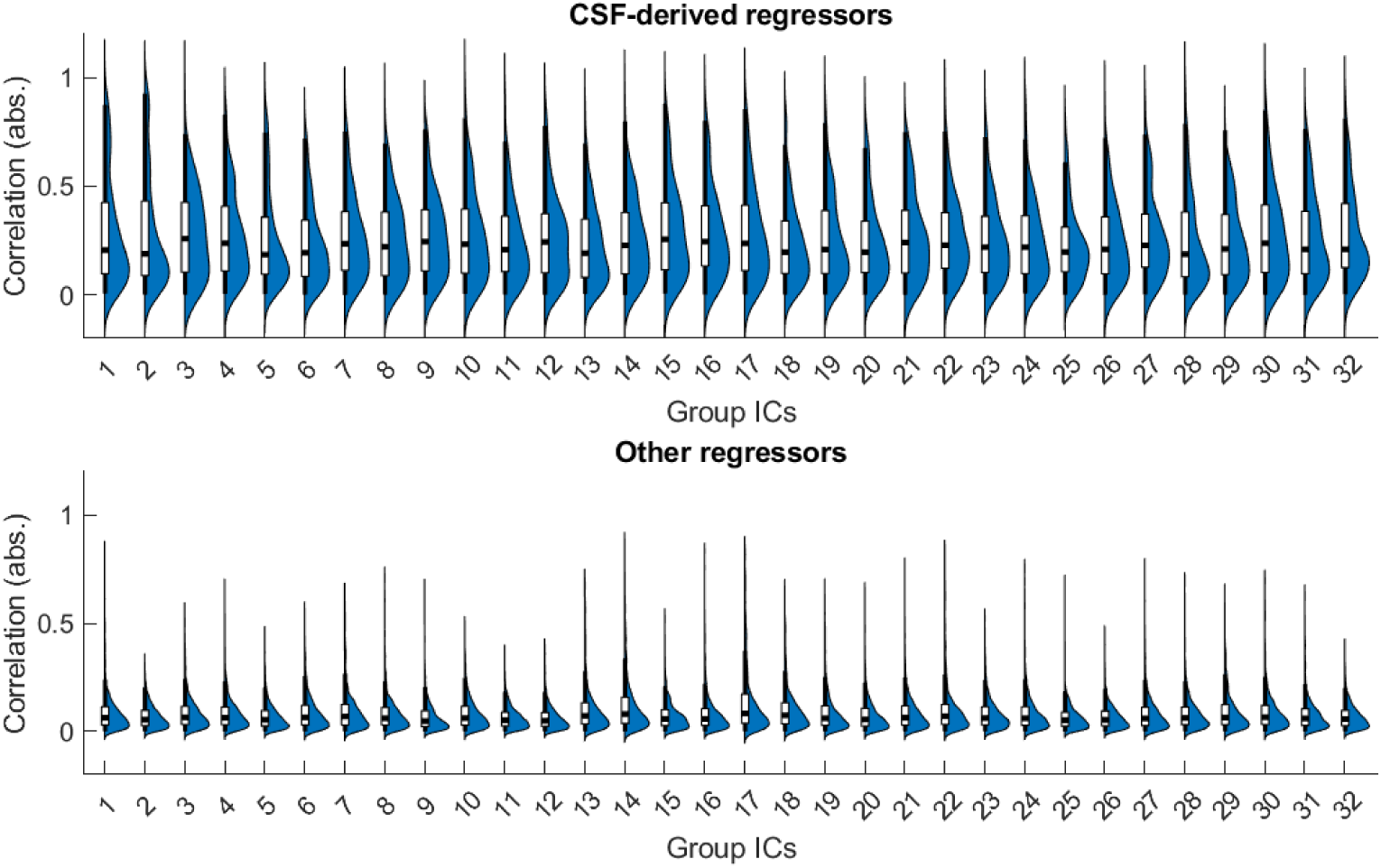
mICs correlate more strongly with CSF-extracted time courses than with other regressors used for denoising. The absolute value of correlation between principal components extracted from CSF near the brainstem and the time-courses of mICs derived from un-denoised-data is higher than that of other regressors used for denoising. The non-neuronal BOLD fluctuations associated with CSF share this increased temporal correlation across all mICs in the group decomposition.

**Figure 11:**
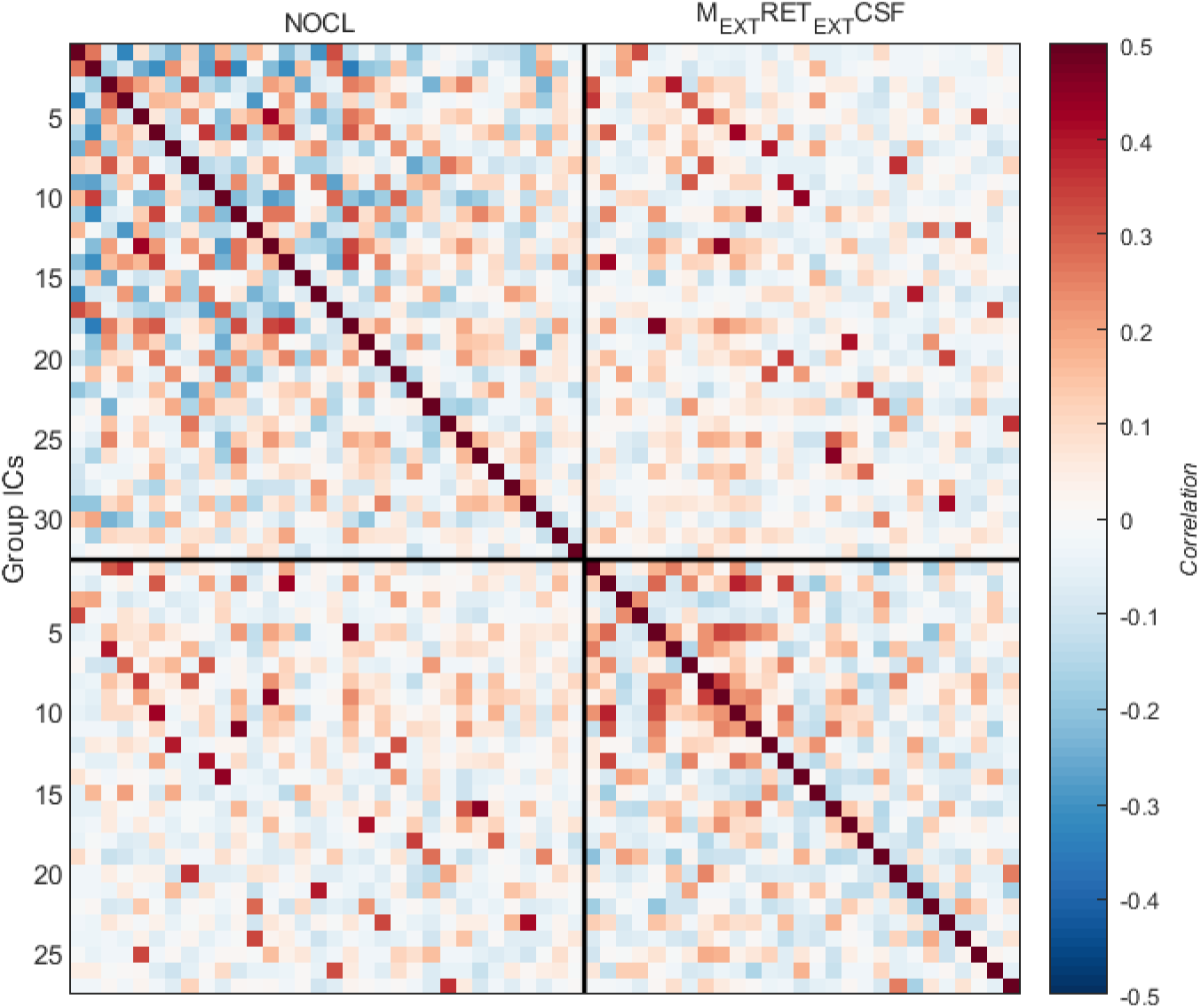
Temporal correlations between the mICs obtained from the un-de-noised (’None’) and most aggressively de-noised (’M_ext_R_ext_CSF’) pipelines show that denoising has a strong effect on the temporal structure of mICA outcomes. As demonstrated in Figure 8, increased de-noising tends to reduce negative correlations between mICs.

Overall, since individual mICs neither replicate the temporal structure of conventional de-noising regressors nor converge to the same temporal structure across de-noising pipelines, we advise that mICA should not be regarded as a substitute for explicit denoising techniques, which in contrast to mICA lend themselves more readily to interpretation. The broad temporal similarities between denoising time-series, particularly CSF, and the mICs obtained through group spatial ICA decomposition suggest that these undesirable artefactual contributions to the BOLD signal tend not to be spatially localized within the brainstem.

We also assessed the spatial replicability of mICs matched across all pipelines. Figure 12 shows that the application of denoising had the greatest effect on spatial characteristics of mICs when CSF-extracted regressors were included in nuisance regression, a pattern reinforced by the observation of split-half reproducibility differentiation for these pipelines at higher dimensionalities (Figure 3).

**Figure 12:**
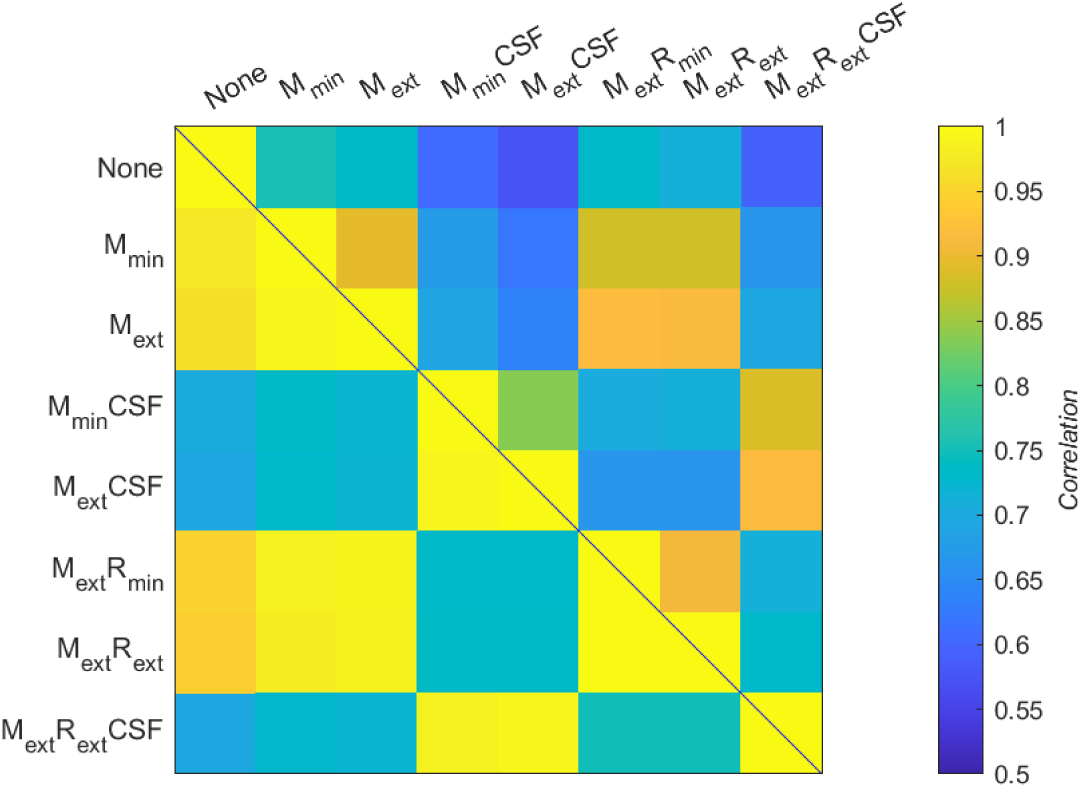
Inter-pipeline spatial reproducibility scores suggest that the effects of CSF-denoising cause the greatest variability between pipelines. Here, the spatial correlation is averaged over all components matched within pairs of pipelines. All pipelines achieve a spatially reproducible (>0.5) score. Upper triangle: median dimensionality implementations; lower triangle: reproducible dimensionality implementations.

### Comparison with low frequency physiological fluctuations

Temporally, only a small proportion of mICs within each pipeline shared characteristics with low frequency physiological time series linked to autonomic function and arousal (heart rate variability and changes in respiration volume per time convolved with physiological response functions). For both dimensionality implementations of group mICA, more extensive physiological de-noising tended to reduce correlation with HRV but not RVT. In fact, the inclusion of CSF de-noising tended to increase the number of components correlated with RVT. Figure 13 shows results for a representative selection of pipelines, while overall results are summarized in Table 3. As expected, the higher dimensionality implementation of mICA yielded more components which were temporally correlated to these physiological sources of variance, though the statistical necessity of correcting for multiple correlations in fact reduced the proportion of components for which such correlations were statistically significant, as the increased dimensionality did not significantly increase the correlation strength for these components.

**Figure 13:**
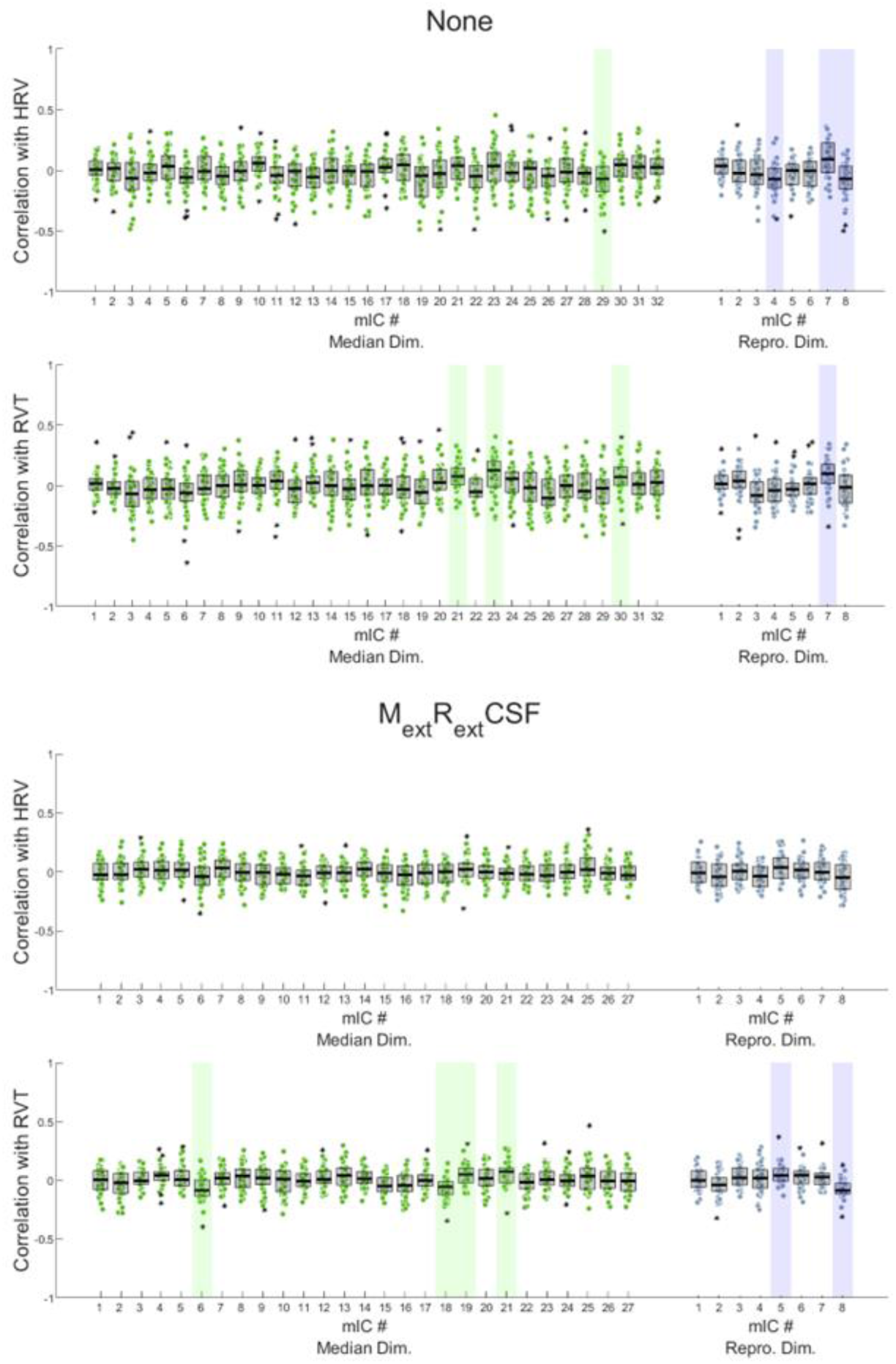
Temporal correlations with respiration volume per time (RVT) are preserved in mICs after denoising, while those with heart rate variability (HRV) are reduced. Shaded regions indicate significant correlation scores (p<0.05; Bonferroni corrected within pipelines).

**Table 3:**
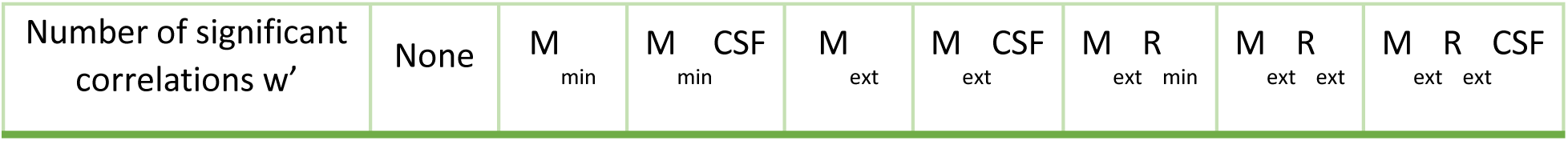
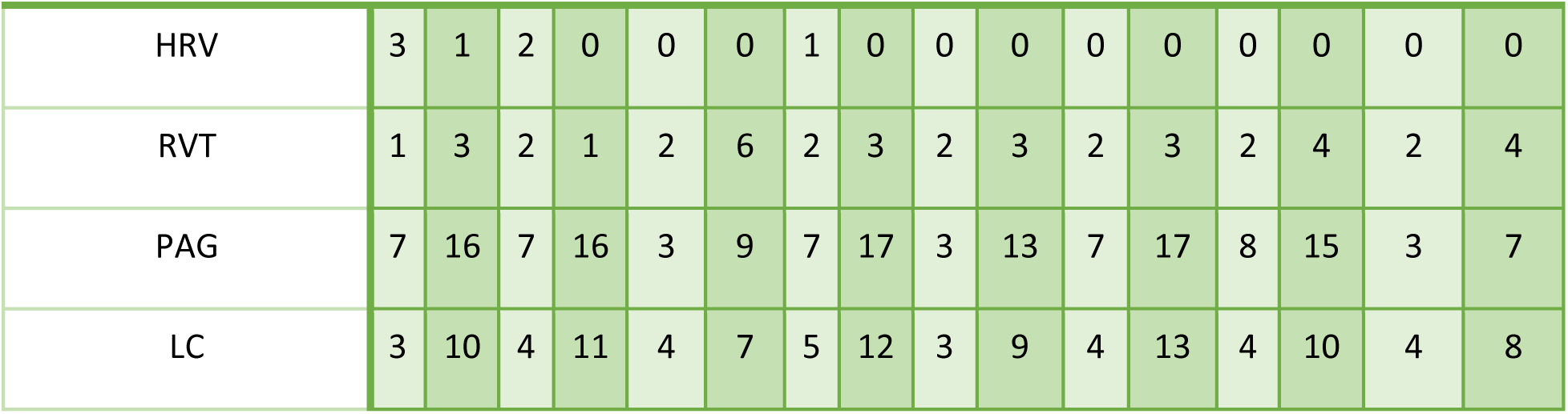
Number of mIC components significantly correlated (p<0.05) with physiological and atlas-extracted time courses. Light shading indicates results for reproducible dimensionality (8) while darker shading represents results for median dimensionality (various across pipelines).

**Table 4:**
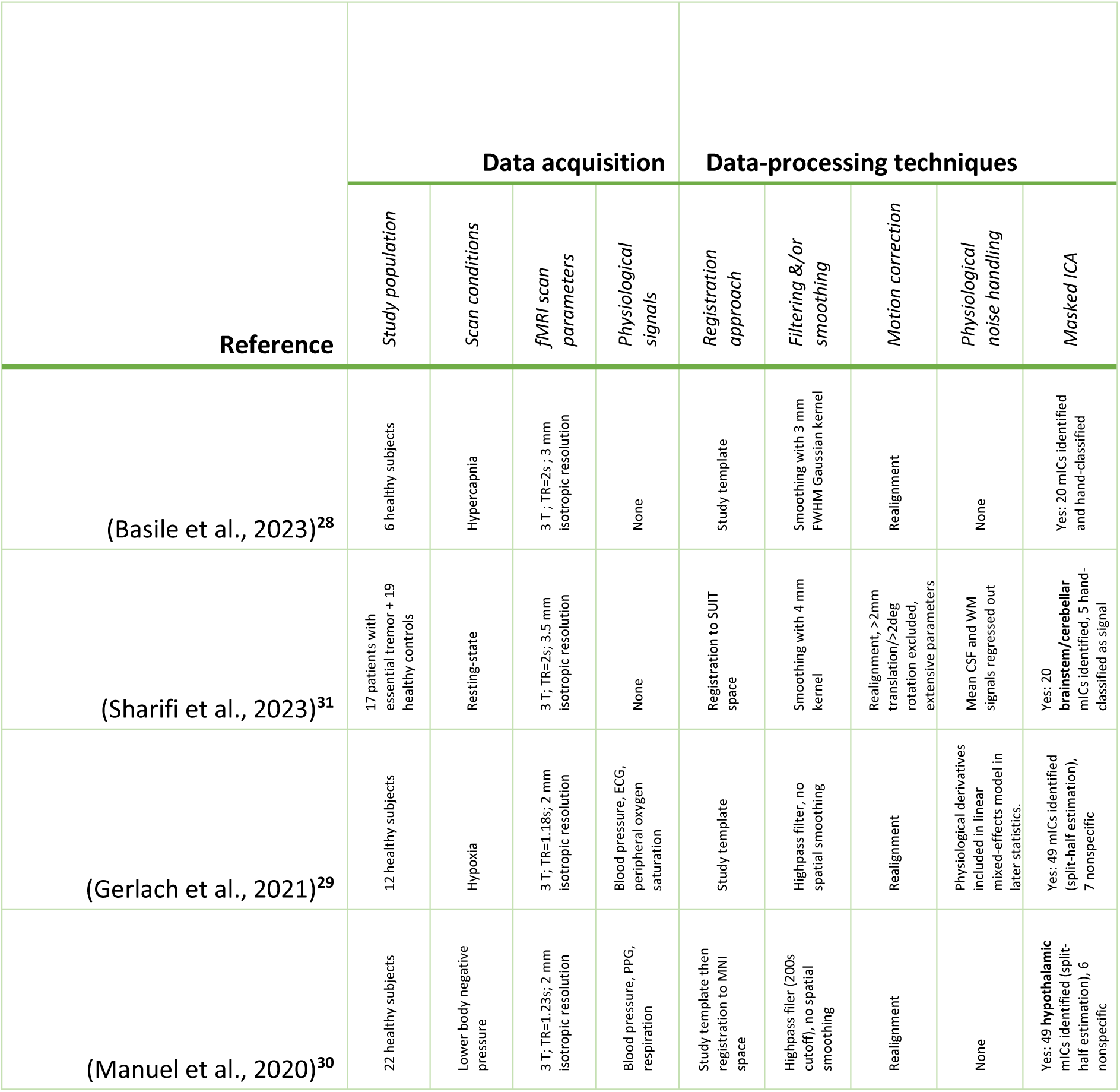
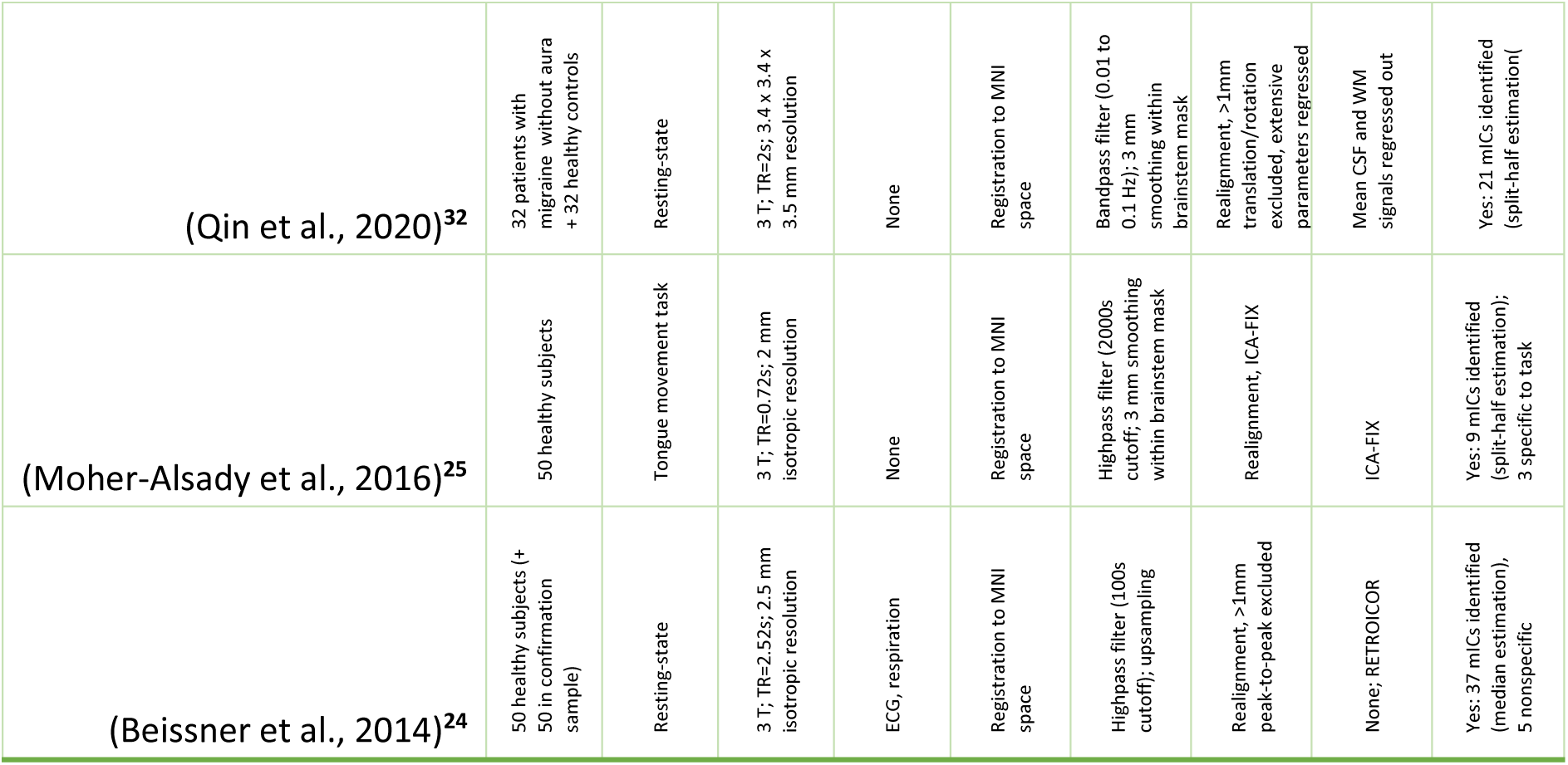
Overview of fMRI studies implementing masked ICA for the brainstem, cerebellum, or hypothalamus. To the authors’ current knowledge, this table presents a comprehensive list of the current studies which have used mICA for subcortical regions including or relevant to the brainstem. Sharifi et al.’s study^31^ of cerebellar connectivity was included as masked ICA was applied to the brainstem and cerebellum jointly. Manuel et al.’s study^30^ of cardiovascular regulation in the hypothalamus was included for its relevance to noise handling during an autonomic task and explicit consideration of connectivity to the brainstem.

### Comparison with atlas-based signal extraction

Examination of whether mICA could replicate atlas-based signal extraction from autonomically relevant brainstem regions such as the locus coeruleus (LC) and periaqueductal gray (PAG) revealed that the higher dimensionality implementation of group mICA yielded a lower proportion of components sharing signal with these regions, which could potentially be advantageous for de-noising using mICs. More extensive physiological denoising tended to reduce correlation with the PAG but not the LC, perhaps because of the latter’s more central location shielding it from physiological noise sources more prominent at the brainstem’s edges. Figure 14 shows results for a representative selection of pipelines, while overall results are summarized in Table 3.

**Figure 14:**
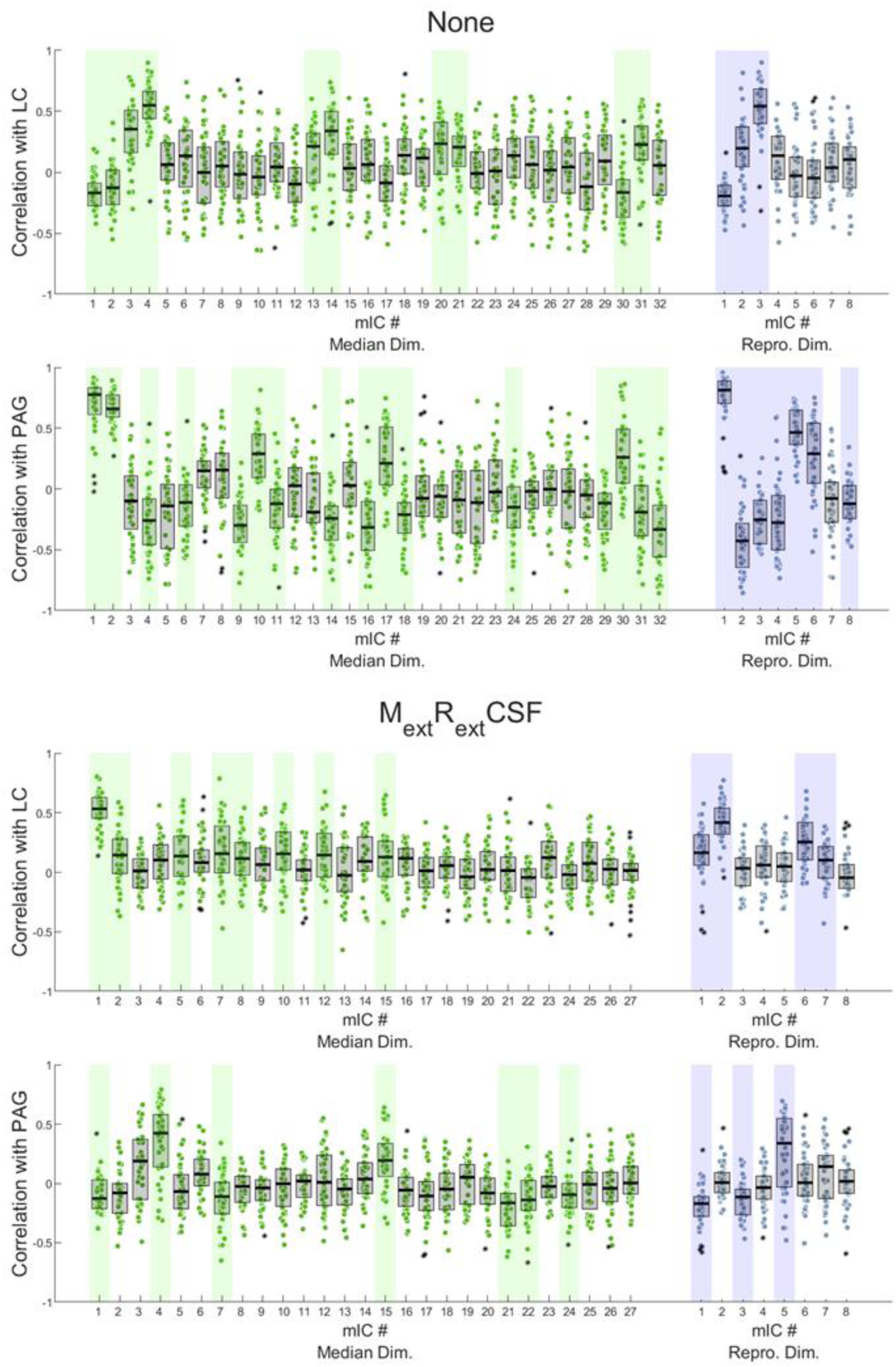
Temporal correlations with signals extracted from the LC and PAG are widely distributed across mICs, although for higher dimensionality implementations (left) this effect is somewhat reduced. Shaded regions indicate significant correlation scores (p<0.05; Bonferroni corrected within pipelines).

## Discussion

### Summary of main findings

We applied mICA across a range of dimensionality selections and preprocessing pipelines and established normative spatial and temporal characteristics of mICs across implementations (Figure 4, Figure 5, Figure 8). We reproduced the majority (20/37) of components spatially identified by Beissner et al. for one or more de-noising pipelines (Figure 7) while observing a higher proportion of noise-aligned components in our dataset. However, for the first time, we show that mICA is temporally weakly correlated to other commonly applied models of motion and physiological noise processes (Figure 9). Moreover, we show that the effects of CSF-based denoising are particularly impactful on mICA outcomes both temporally (Figure 10) and spatially (Figure 12). Evaluating classification techniques, we demonstrated general agreement between specificity values and hand-labelling outcomes for mICs (Figure 6), but observed that some mICs with noise-like spatial distributions achieve high specificity scores and require a more nuanced approach for delineation. Lastly, we made a comparison of mIC time courses with slow physiological fluctuations and atlas-extracted signals to highlight their respective overlap in accounting for temporal variance within the brainstem (Figure 13, Figure 14).

Overall, our analyses suggest that the effects of motion and physiological noise permeate even a conservative masking of brainstem data at the group level. These strong physiological noise sources include but are not limited to bulk head motion, pulsatility-related cardiac motion, magnetic susceptibility changes stemming from respiration, vasodilation varying with blood gas levels during respiration, and interactions between cardiac and respiratory processes^14,52,53^. Within the brainstem itself, large BOLD signal variance reflects not only the presence of strong physiological noise sources, but is also potentially attributable to a significant role of low-frequency functional connectivity^54^. The application of mICA continues to hold promise for a spatial decomposition of noise and signal effects in the brainstem. However, our results demonstrate that mICA should not be seen as a substitute for temporal denoising techniques.

### Effects of preprocessing decisions on mICA outcomes

We have demonstrated that CSF-denoising particularly affects the spatial and temporal characteristics of mICA-identified sources. However, all denoising pipelines were found to differentially affect the temporal interpretation of mICs, both through the temporal dynamics of the mICs themselves, and their shared temporal variance with whole-brain data. Since mICs are typically used in conjunction with whole-brain data to establish the connectivity of brainstem sources (Table 4), merely discarding mICs classified as noise from further analysis is insufficient to account for noise both within and exterior to the brainstem. This is particularly true in the context of autonomic investigations, where estimations of spontaneous whole-brain connectivity must be carefully distinguished from global physiological noise^55^.

By providing an overview of the effects of various denoising techniques, we hope that this study will motivate the incorporation of dedicated and specific denoising in conjunction with mICA to improve the quality of their source identification and whole-brain statistical analyses.

### Classification of mICA components

Although mICA components were originally proposed while accompanied by robust reporting of spatial maps, reproducibility, and specificity scores^24^, subsequent implementations of mICA^29,30,32^ have relied on specificity or specificity-adjacent metrics to classify mICs without explicitly discussing their underlying characteristics. Moreover, in the two most recent applications of mICA at the time of the present study, the use of specificity or correlation with an external task to identify mICs as signal has given way to hand classification^28,31^ (Table 4). We have shown that neither approach provides a completely objective classification criterion: components hand-labelled as noise may have high specificity (Figure 6), while hand-labelling group-level components may not be equipped to distinguish ambiguous components (Figure 5). Clear reporting of classification criteria is essential for beginning to assess the utility of mICA’s application, particularly in resting-state data. We suggest that going forward, beyond specificity scores, whole-brain connectivity maps may be leveraged during hand-labelling for a more nuanced interpretation of noise signatures which are well-characterized in a whole-brain context.^48^

It should also be noted that while our study explored the implementation of mICA on a 7T EPI dataset optimized to provide a high spatial resolution in the brainstem, other tailored data acquisitions could offer greater versatility in approaches to classification. For example, growing interest in multi-echo fMRI sequences for brainstem imaging^56,57^ have yet to leverage robust TE-dependent ICA classification techniques^58^ in a masked context. Moreover, ultrafast sequences could better leverage the temporal dimension of fMRI data with masked ICA to better distinguish global noise signatures.^59^

### Dimensionality selection and the identification of brainstem nuclei

As a further point regarding the implementation of resting-state mICA, we note that a reproducibility-based approach to dimensionality selection may not be granular enough to spatially isolate signal from noise (Table 3). However, determining the number of independent sources which can be meaningfully identified in the brainstem using resting-state fMRI, whether at the individual or group level, is not trivial. While we showed that increasing the dimensionality of mICA decreased the average spatial reproducibility within a split-half comparison of the group (Figure 3) and chose to alternatively implement higher-valued median-estimated dimensionalities (ranging from 27-32) contrary to current mICA guidelines^25^, it may be the case that even higher dimensionality decompositions with mICA would improve the identification of brainstem nuclei. Recent studies have suggested applying segmentations of between and 58^60,61^ and 84^62^ functionally differentiable sub-regions within the brainstem at rest, even for lower resolution acquisitions.

Given the significant differences in study designs, dataset compositions, preprocessing approaches, and selection of mICs to include in further analysis which currently exist in papers published which use mICA in the brainstem (Table 4), further evidence is warranted to substantiate the claim that mICA can be used to measure the activation of brainstem nuclei in resting-state data. Deeper investigation into the correspondence between atlas-based signal extraction of autonomic and other nuclei with mICA sources within the brainstem would help to build a case for the relevance of mICA-based studies to the broader brainstem literature; the use of autonomic task data may likewise present a stronger argument for mICA sensitivity to brainstem activations in the future. Nonetheless, based on the outcomes of this study, no implementation of mICA on our resting-state data was able to identify signal fluctuations linked to regulation of physiological processes by autonomic nuclei in the brainstem.

It should be noted that one promising avenue to identify brainstem regions implicated in autonomic regulation leverages whole-brain data-driven approaches to increase sensitivity to relevant brainstem activation. A recent study reported twelve regions-of-interest in the brainstem that overlapped with whole-brain ICA networks involved in autonomic control.^63^ It is thus possible that, despite strong physiological noise in the brainstem, whole-brain data-driven analyses may offer a worthwhile alternative pathway to disentangling signal from noise in autonomic contexts.

### Limitations of the present approach

We designed a suite of preprocessing pipelines and mICA implementation which adhered closely to those used previously in mICA benchmarking papers^24,25^ to better address questions of replicability. However, our results demonstrate that the central premise of mICA – that is, masking the brainstem to avoid contamination by exterior sources of noise – does not entirely succeed while following this implementation. Indeed, we identified a higher proportion of noise sources than previously reported (Table 3).

While diverging from previous mICA implementations, our group-level analysis could have been improved in several regards: namely, in mask delineation and registration to a common space. In particular, registration using tools designed explicitly to achieve accurate subcortical alignment^64^ could make a significant difference in group-level analyses going forward. Beyond group-level mICA in which inter-subject anatomical variability and registration inaccuracies might give rise to the presence of noise signatures within the mask, individual mICA applied within each subject’s functional space could offer an alternative. For denoising, a single-subject decomposition would provide a more ready comparison with whole brain ICA applications.^48^ Such an approach has been previously demonstrated for mapping the LC in individual fMRI scans^34^ with considerable variability across subjects.

Although we attempted to explore the implementation of single-subject mICA, we encountered several problems leading to this investigation’s omission from the current study. First, accurate automated segmentation of subject-specific brainstem masks proved challenging in our structural dataset. Second, the estimation of the dimensionality of the number of underlying sources in individual functional space was highly inconsistent depending on the estimation technique and amount of smoothing. Since this technique did not allow a reproducibility-based approach to dimensionality estimation – and indeed, the number of sources for each subject might be expected to vary significantly based on the intensity of motion or other physiological changes during their scan – we could not identify a robust way to proceed within the scope of this paper.

## Conclusions

We have evaluated the outcomes of de-noising and dimensionality selection on the application of mICA to brainstem resting-state data. Overall, regardless of preprocessing approach, we do not replicate the high specificity values previously obtained for resting-state data or fully replicate previously reported spatial sources in 7T resting-state data.^24^ We also observe that mICA does not replicate the extraction of time courses from an established atlas of brainstem nuclei^51^, although the accuracy of these nucleic segmentations may be limited by the accuracy of subcortical registration.^26^ Moreover, we observe fairly consistent behaviour of mICs across de-noising pipelines and weak correlations with slow physiological effects encapsulated by HRV and RVT, suggesting that mICA should not be regarded as equivalent to or substituted for widely-used physiological denoising techniques. In general, we advocate for close inspection and thorough reporting of mIC decomposition, as the use of automated classification schemes based on specificity such as those often used with task data^25,29,30^ do not adequately distinguish noise characteristics of these data.

## Data and code availability

For the ongoing availability of the raw data used in this study, please refer to its original publication^35^. Analyzed data at the group level (including mICA outcomes) are available from the corresponding author upon reasonable request. Code used for this study is available from the corresponding author upon reasonable request.

## Author contributions

Mary Miedema: conceptualization; formal analysis; methodology; visualization; writing—original draft preparation; writing—review & editing. Kyle Pattinson: data acquisition; writing—review & editing. Georgios Mitsis: conceptualization; supervision; writing—review & editing.

## Acknowledgements

The authors wish to thank Olivia Faull Harrison for her role in the original data acquisition which was shared with this study.

