## Supplemental Figures for "Towards the implementation and interpretation of masked ICA for identifying signatures of autonomic activation in the brainstem with resting-state BOLD fMRI"

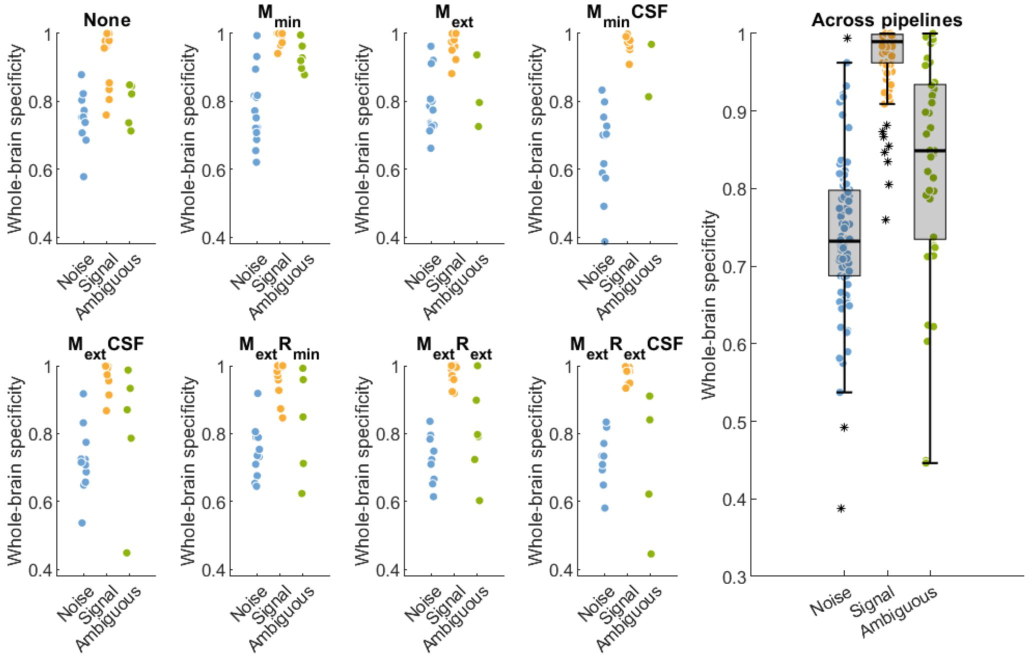


*Figure 1: Hand-labelling of individual components with corresponding specificity values (calculated using smoothed whole-brain data). Results for each individual pipeline (median dimensionality) are shown on the left; their aggregate is on the right.*

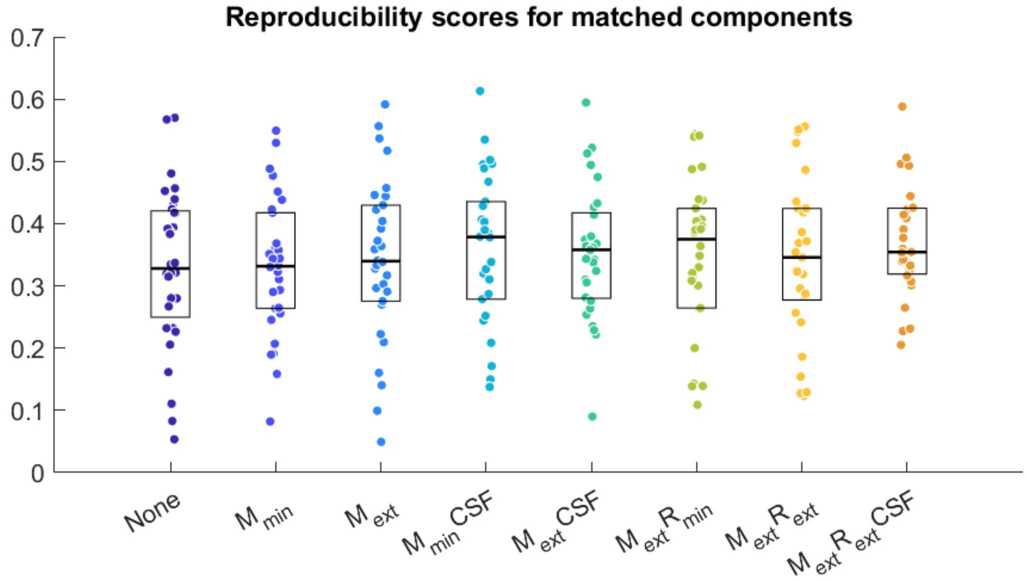


*Figure 2: The distribution of spatial reproducibility scores of mICs matched with Beissner et al.'s (2014) originally reported 37 resting-state mICs is not significantly improved by denoising.*

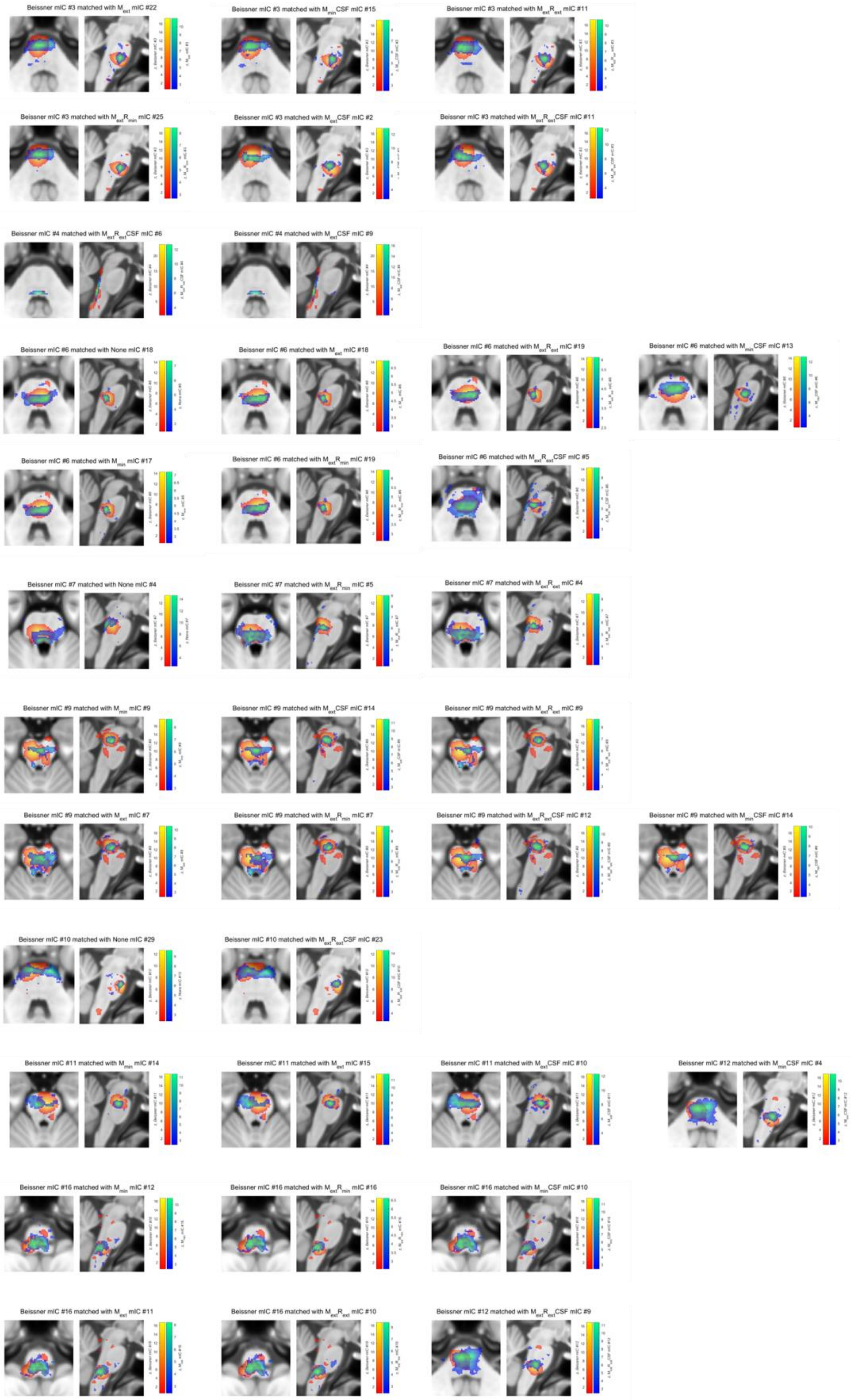

Beissner mIC #3 matched with $M_{ext}$ mIC #22
Beissner mIC #3 matched with $M_{min}$CSF mIC #15
Beissner mIC #3 matched with $M_{ext}R_{ext}$ mIC #11
Beissner mIC #3 matched with $M_{ext}R_{min}$ mIC #25
Beissner mIC #3 matched with $M_{ext}$CSF mIC #2
Beissner mIC #3 matched with $M_{ext}R_{ext}$CSF mIC #11
Beissner mIC #4 matched with $M_{ext}R_{ext}$CSF mIC #6
Beissner mIC #4 matched with $M_{ext}$CSF mIC #9
Beissner mIC #6 matched with None mIC #18
Beissner mIC #6 matched with $M_{ext}$ mIC #18
Beissner mIC #6 matched with $M_{ext}R_{ext}$ mIC #19
Beissner mIC #6 matched with $M_{min}$CSF mIC #13
Beissner mIC #6 matched with $M_{min}$ mIC #17
Beissner mIC #6 matched with $M_{ext}R_{min}$ mIC #19
Beissner mIC #6 matched with $M_{ext}R_{ext}$CSF mIC #5
Beissner mIC #7 matched with None mIC #4
Beissner mIC #7 matched with $M_{ext}R_{min}$ mIC #5
Beissner mIC #7 matched with $M_{ext}R_{ext}$ mIC #4
Beissner mIC #9 matched with $M_{min}$ mIC #9
Beissner mIC #9 matched with $M_{ext}$CSF mIC #14
Beissner mIC #9 matched with $M_{ext}R_{ext}$ mIC #9
Beissner mIC #9 matched with $M_{ext}$ mIC #7
Beissner mIC #9 matched with $M_{ext}R_{min}$ mIC #7
Beissner mIC #9 matched with $M_{ext}R_{ext}$CSF mIC #12
Beissner mIC #9 matched with $M_{min}$CSF mIC #14
Beissner mIC #10 matched with None mIC #29
Beissner mIC #10 matched with $M_{ext}R_{ext}$CSF mIC #23
Beissner mIC #11 matched with $M_{min}$ mIC #14
Beissner mIC #11 matched with $M_{ext}$ mIC #15
Beissner mIC #11 matched with $M_{ext}$CSF mIC #10
Beissner mIC #12 matched with $M_{min}$CSF mIC #4
Beissner mIC #16 matched with $M_{min}$ mIC #12
Beissner mIC #16 matched with $M_{ext}R_{min}$ mIC #16
Beissner mIC #16 matched with $M_{min}$CSF mIC #10
Beissner mIC #16 matched with $M_{ext}$ mIC #11
Beissner mIC #16 matched with $M_{ext}R_{ext}$ mIC #10
Beissner mIC #12 matched with $M_{ext}R_{ext}$CSF mIC #9

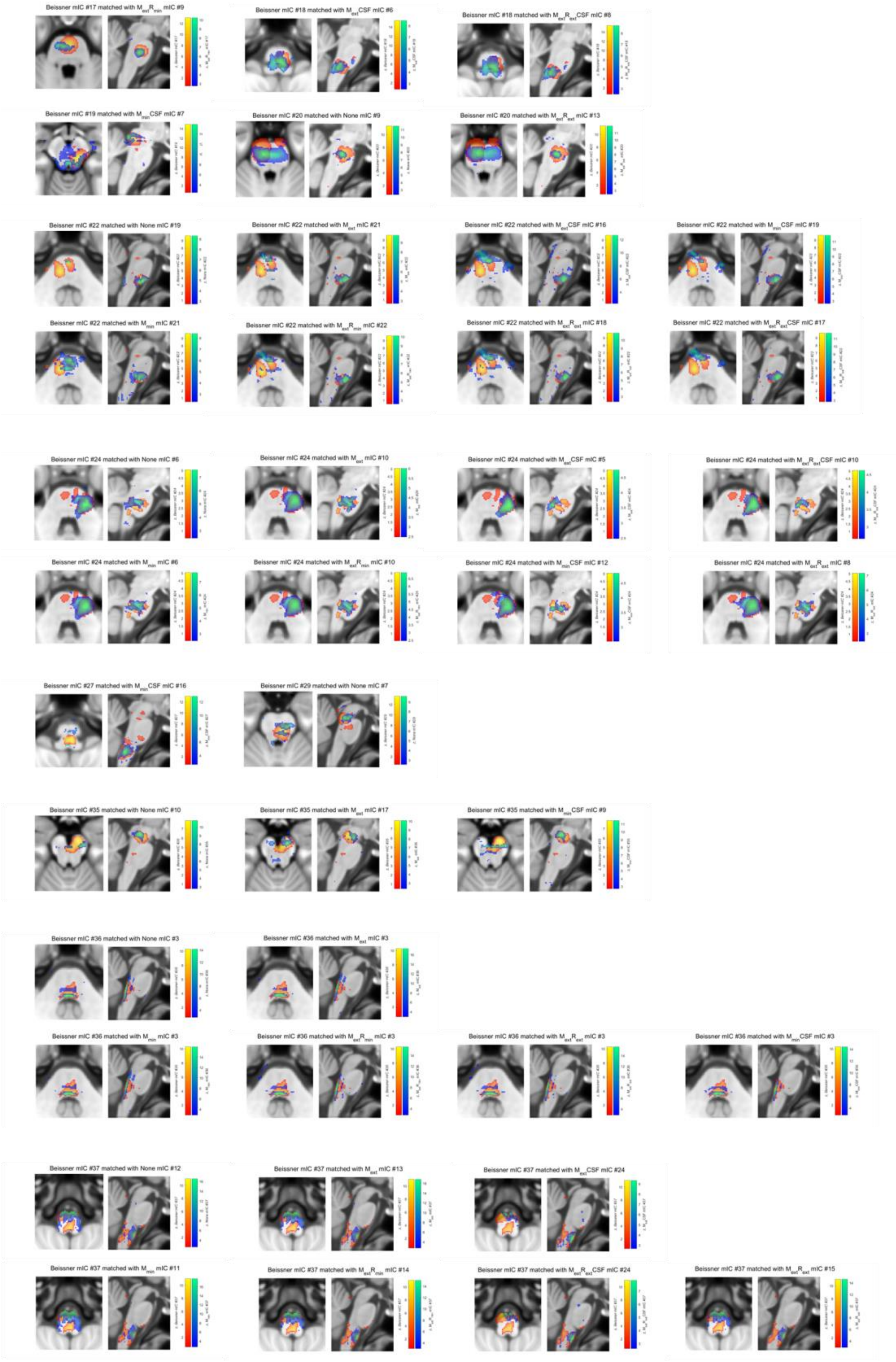


*Figure 3: Matched reproducible (spatial correlation > 0.4) components from this study and Beissner et al.'s original masked ICA paper (2014) from all denoising pipelines (median dimensionalities).*

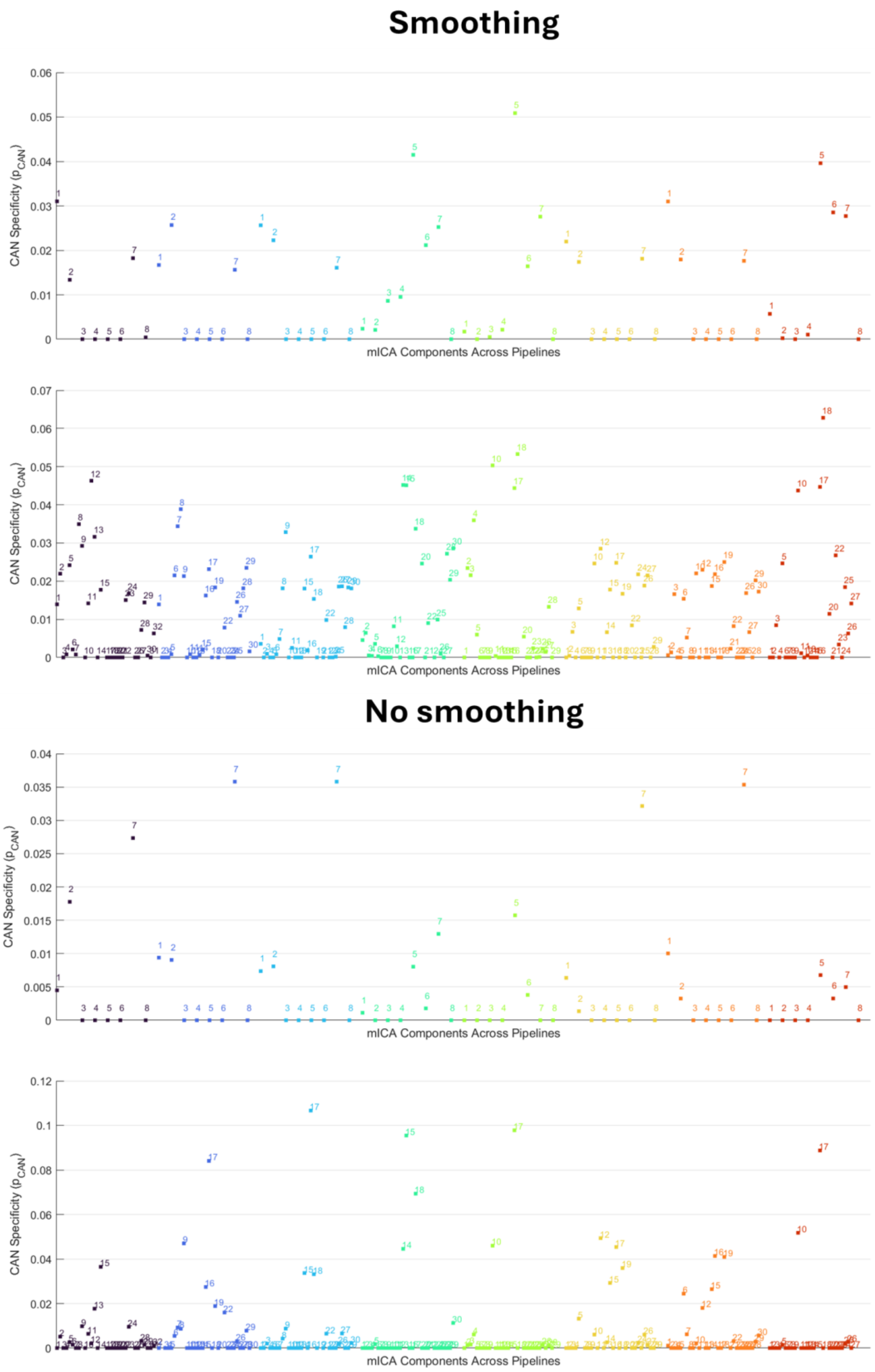


*Figure 4: CAN specificity scores across pipelines were very low for all components, although higher values were achieved for the more granular median dimensionality application of mICA (bottom panels).*
